# Structure and replication cycle of a virus infecting climate-modulating alga *Emiliania huxleyi*

**DOI:** 10.1101/2023.06.30.547180

**Authors:** Miroslav Homola, Carina R. Büttner, Tibor Füzik, Pavel Křepelka, Radka Holbová, Jiří Nováček, Marten Chaillet, Friedrich Förster, William H. Wilson, Declan C. Schroeder, Pavel Plevka

## Abstract

The globally distributed marine alga *Emiliania huxleyi* produces reflective calcite disks (coccoliths) that increase the albedo of ocean water and thus reduce the heat absorption in the ocean, which cools the Earth’s climate. The population density of *E. huxleyi* is restricted by nucleocytoplasmic large DNA viruses, including *E. huxleyi* virus 201 (EhV-201). Despite the impact of *E. huxleyi* viruses on the climate, there is limited information about their structure and replication. Here we show that the dsDNA genome inside the EhV-201 virion is protected by an inner membrane, capsid, and outer membrane decorated with numerous transmembrane proteins. The virions are prone to deformation, and parts of their capsids deviate from the icosahedral arrangement. EhV-201 virions infect *E. huxleyi* by using their fivefold vertex to bind to a host cell and fuse the virus’s inner membrane with the plasma membrane. Whereas the replication of EhV-201 probably occurs in the nucleus, virions assemble in the cytoplasm at the surface of endoplasmic reticulum-derived membrane segments. Genome packaging initiates synchronously with the capsid assembly and completes through an aperture in the forming capsid. Upon the completion of genome packaging, the capsids change conformation, which enables them to acquire an outer membrane by budding into intracellular vesicles. EhV-201 infection induces a loss of surface protective layers from *E. huxleyi* cells, which allows the continuous release of virions by exocytosis. Our results provide insight into how EhVs bypass the surface protective layers of *E. huxleyi* and exploit the organelles of an infected cell for progeny assembly.

## Introduction

*Emiliania huxleyi* is a globally distributed single-celled marine alga known for its ability to multiply quickly in large ocean areas, resulting in blooms covering hundreds of thousands of square kilometers (*1–3*). Spherical *E. huxleyi* cells, with a diameter of 4-5 µm, are protected by calcite disks called coccoliths, which increase the albedo of seawater by reflecting light, and thus decrease the amount of heat from solar radiation absorbed by oceans (*2, 4*). This alga is an important component of the global carbon cycle, as the coccoliths shed from the cells descend to the sea bottom and serve as a sink for carbon dioxide (*5*). Furthermore, dimethyl sulfide and other compounds released by *E. huxleyi* promote the condensation of atmospheric aerosol droplets and the formation of clouds that reflect sunlight (*6*). These properties, in combination with broad distribution and high abundance, enable *E. huxleyi* to exert a cooling effect on the Earth’s climate (*2, 7, 8*).

*E. huxleyi* is susceptible to infection by nucleocytoplasmic large DNA viruses (NCLDVs), which reduce the population density of the alga and alter its impact on the climate (*9–11*). NCLDVs infecting algae belong to the family *Phycodnaviridae* from the order *Algavirales* (*12, 13*). The most extensively studied representative of algal viruses is Paramecium bursaria chlorella virus 1 (PBCV-1) from the genus *Chlorovirus* (*14–16*). Emiliania huxleyi virus 201 (EhV-201) and closely related EhV-86 belong to the genus *Coccolithovirus* (*17–19*). More than twenty viruses from the family *Phycodnaviridae* that infect *E. huxleyi* have been isolated; however, only EhV-86 genome has been fully sequenced to date (*17, 20*). The genome of EhV-86 has a size of 407 kbp with 472 predicted protein-coding sequences (*17*).

Virions of coccolithoviruses are unique within the *Phycodnaviridae* family, because they contain not only a membrane inside the capsid, but also an additional membrane wrapped around the outer capsid surface (*21, 22*). It has been speculated that EhV-86 delivers its genome into cells by the fusion of its outer membrane with a cell membrane, similar to enveloped animal viruses (*22*). Furthermore, it has been proposed that the capsid of EhV-86 enters the cytoplasm intact and releases its genome into the nucleus (*22*). EhV-86, similar to other phycodnaviruses, probably replicates its genome in the cell nucleus, but progeny particles assemble in the cytoplasm (*22*). Towards the end of the virus replication cycle, which takes approximately five hours, the cytoplasm of *E. huxleyi* cells contains tens of progeny virions (*22*). It has been speculated that EhVs acquire their outer membrane by budding from the host cell membrane (*21, 22*). Despite previous studies of EhV-86 and possible analogies with the better-studied PBCV-1 (*14–16*), many aspects of EhV structure and replication remain unknown.

Here we used various electron microscopy approaches to show that the EhV-201 infection process is different than previously inferred based on data obtained using lower resolution techniques. The EhV-201 virion delivers its genome into the algal cytoplasm by fusing its inner membrane with the plasma membrane. The capsid, together with the outer membrane, remain attached to the cell surface. After genome replication in the nucleus, EhV-201 capsid assembly initiates in the cytoplasm synchronously with genome packaging on membrane segments derived from the endoplasmic reticulum. Upon completion of the genome packaging, EhV-201 particles bud into intracellular vesicles and thus acquire their outer membrane. EhV-201 infection induces the loss of surface protective layers from *E. huxleyi* cells, which enables the continuous release of progeny virions by exocytosis.

## Results & Discussion

### Structure of EhV-201 virion

Virions of EhV-201 are spherical in shape with a maximum outer diameter of 211 nm (Fig. 1A, S1). The EhV-201 genome is protected by a 4.2-nm-thick inner membrane, a 6.1-nm-thick capsid, and a 6.1-nm-thick outer membrane (Fig. 1A-C, S2). Unlike virions of NCLDVs with isometric capsids which have been structurally characterized to date (*14, 16, 23–26*), those of EhV-201 are deformed, and parts of their capsids lack angular icosahedral features (Fig. 1A, S1). We used sub-tomogram averaging to reconstruct a vertex with regular features and expanded the map according to icosahedral symmetry to obtain a complete EhV-201 virion structure with a resolution of 18 Å (Fig. 1DE, S3A-C, Table S1). The virion reconstruction enabled the identification of three types of transmembrane proteins embedded in the outer membrane: (i) The central vertex proteins located around fivefold symmetry axes bind to pentamers of capsid proteins (Fig. 1DE, S4). The many capsid proteins that form pentamers in capsids of NCLDVs contain large insertions that protrude above the capsid surface (*12*). Therefore, because of the limited resolution of the cryo-EM reconstruction, we cannot exclude the possibility that the inner vertex proteins are domains of the penton proteins of EhV-201. (ii) The peripheral vertex proteins are positioned around the inner vertex proteins, and each of them binds to one hexamer of major capsid proteins surrounding the pentons (Fig. 1DE, S4). (iii) Elongated ridges that cover most of the EhV-201 virion surface are formed by dimers of ridge proteins. Each dimer of ridge proteins binds to two underlying hexamers of major capsid proteins (Fig. 1DE, S4). The EhV-201 virion contains sixty copies of each central and peripheral vertex protein, and at least 1320 copies of ridge protein (Fig. 1E).

**Fig. 1.**
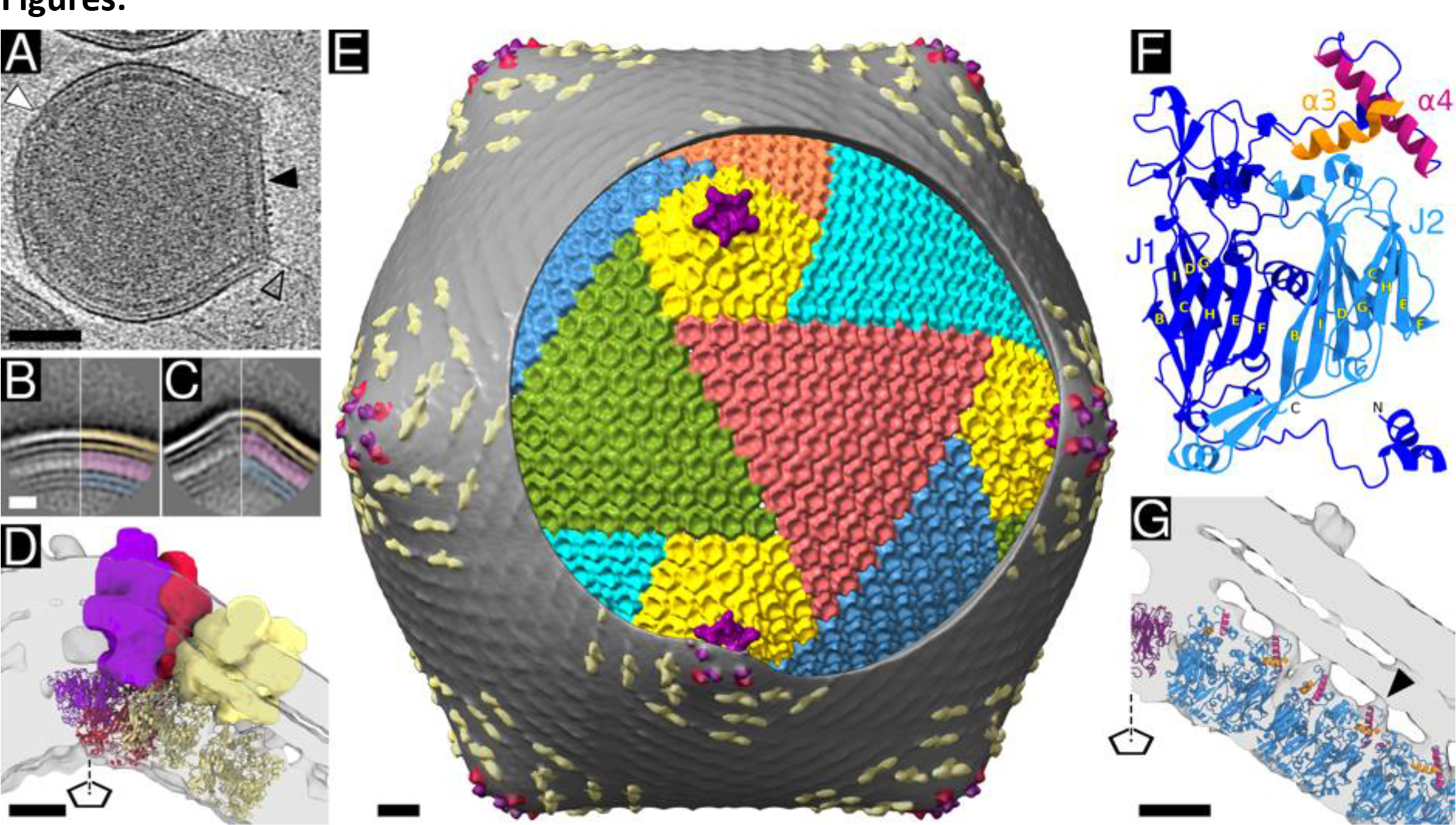
Structure of EhV-201 virion. **(A)** Central section from cryo-tomogram of EhV-201 virion. The part of the particle exhibiting the straight edges and angular vertices that are symptomatic of icosahedral arrangement is indicated by a black arrowhead. The deformed part of the virion is indicated by a white arrowhead. A filament protruding from a virion vertex is indicated by a transparent arrowhead with a black outline. Scale bar 50 nm. **(BC)** Reference-free two-dimensional class averages of round (B) (N = 922) and angular (C) (N = 1,012) segments of EhV-201 virions. Both segments possess identical sequences of surface layers. The layers in the right halves are differentiated: the inner membrane in blue, the capsid in magenta, and the outer membrane in orange. Scale bar 10 nm. **(D)** Cross section of cryo-ET density of angular EhV-201 virion vertex determined to resolution of 13 Å by sub-tomogram averaging. Transmembrane proteins are shown in surface representation and distinguished by colors: the central vertex protein in magenta, peripheral vertex protein in red, and dimer of ridge proteins in light yellow. The capsid proteins are shown in cartoon representation and colored the same as the transmembrane protein they interact with. Models of the capsid proteins were calculated using AlphaFold2 (*27*). The position of the fivefold symmetry axis is indicated by a pentagon and a dashed line. Scale bar 5 nm. **(E)** Composite map of EhV-201 virion. The surface of the virion is covered with the outer membrane (grey) with central (magenta) and peripheral (red) vertex proteins and dimers of ridge proteins (light yellow and grey ripples on the virion surface). A circular region of the outer membrane was removed to reveal the arrangement of the major capsid proteins forming the capsid. Pentamers of capsid proteins are shown in magenta. Pseudo-hexamers of major capsid proteins belonging to the penta-symmetrons are shown in yellow, whereas those forming tri-symmetrons are in various other colors. Scale bar 10 nm. **(F)** AlphaFold2-predicted structure of EhV-201 major capsid protein with double jelly roll fold. The domains J1 and J2 are colored in dark and light blue, respectively. Each domain contains two four-stranded β-sheets with the β-strands conventionally named BIDG and CHEF. Domain J1 contains an insertion between β-strands D and E, which forms amphipathic helices α3 (orange) and α4 (magenta). **(G)** Cross section of cryo-ET density of angular EhV-201 virion vertex showing interactions of amphipathic helices α3 and α4 from J1 domain of major capsid proteins with outer virion membrane (an example is indicated by a black arrowhead). The position of the fivefold symmetry axis is indicated by a pentagon and a dashed line. Scale bar 5 nm.

The capsid of the mature EhV-201 virion has a maximum diameter of 199 nm and a triangulation number of 169 (*h*=7, *k*=8) (Fig. 1E, S5). Capsomers of major capsid proteins are organized into penta-symmetrons and tri-symmetrons, as is in other NCLDVs (Fig. 1E). The structure of the EhV-201 major capsid protein, predicted using AlphaFold2 (*27*), has the characteristic double jellyroll fold of the capsid proteins of NCLDVs and other viruses (Fig. 1F) (*28*). The jellyroll cores J1 and J2 are each composed of two β-sheets named according to the convention BIDG and CHEF (Fig. 1F) (*28*). Three copies of the major capsid protein form a capsomer with quasi-sixfold symmetry (Fig. S6).

The EhV-201 inner virion membrane is less well resolved than the outer one, indicating higher variability in its structure between individual particles (Fig. 1A-C, S7). The reconstruction provides no indication of minor capsid proteins mediating contacts between the inner membrane and the capsid. This finding is in contrast with NCLDVs, such as Tokyovirus and PBCV-1, whose inner membranes are stabilized by interactions with capsid proteins (*14, 16, 29*).

### Interactions between the outer membrane and capsid

The transmembrane proteins from the EhV-201 outer membrane bind to the capsid; however, there are additional interactions between the major capsid proteins and the outer membrane. The major capsid protein of EhV-201 contains a 96-residue-long loop between β-strands D and E of the J1 jellyroll domain (Fig. 1F, S6). The loop is predicted to form helices α3 and α4, which are 13 and 20 residues long, respectively, and are positioned at the outer surface of the capsid (Fig. 1FG, S7). Helices α3 and α4 contain hydrophobic residues organized in an amphipathic α-helical arrangement, which pre-disposes them to bind to membranes (Fig. S8). Fitting the predicted EhV-201 capsomer structure into the sub-tomogram reconstruction of the virion vertex positions helices α3 and α4 inside densities connecting the capsid to the outer membrane (Fig. 1G). Therefore, we speculate that the amphipathic helices stabilize the attachment of the outer virion membrane to the capsid.

The abundant capsid-outer membrane contacts may enable EhV-201 virions to withstand deformation without negatively affecting the infectivity of the virus. A comparison of the sequences of major capsid proteins of NCLDVs indicates that the amphipathic helices α3 and α4 are a unique feature of coccolithoviruses among the viruses from the family *Phycodnaviridae* (Fig. S9).

### Filaments attached to vertices of EhV-201 virions

Tomograms of 96% of EhV-201 virions (N = 50) contained at least one 3.3 nm thick (SD = 0.5, N = 20) and 30 - 150 nm long (mean = 72, SD = 31, N = 20) filament protruding from a fivefold particle vertex (Fig. 1A, S10). The filaments emerge from the outer membrane, but their exclusive positioning at the vertices provides evidence that they bind to specific sites at the capsid (Fig. 1A, S10). We identified particles containing more than one such filament, indicating that it is unlikely that the filament is a feature of a special vertex in the EhV-201 virion (Fig. S10). The classification of sub-tomograms of EhV-201 virion vertices did not identify a sub-population of vertices containing the putative filament. The filament may be a feature that is too weak and flexible to be detected by the classification procedure. The putative function of EhV-201 filaments in *E. huxleyi* cell infection may be similar to that of PBCV-1, which has been observed to attach to cell walls via hair-like fibers (*30*). Furthermore, there is evidence of specific protein receptor-ligand interactions during initial EhV attachment engagement, which may be mediated by the putative fibers (*31*).

### EhV-201 attachment and genome delivery

Many NCLDVs characterized to date deliver their genomes into the host cell cytoplasm by fusing their inner capsid membrane with the host plasma membrane (*14, 32–35*). This mechanism of genome delivery is characterized by the capsid and emptied inner membrane sack remaining attached to the surface of an infected cell (*14, 34*). In contrast, it has been speculated that EhV-86 infects cells via the fusion of its outer virion membrane with the plasma or endosome membrane, which would result in the delivery of the genome enclosed within the inner membrane and capsid into the host cytoplasm (*21*).

We used serial block-face scanning electron microscopy of vitrified and resin-embedded *E. huxleyi* cells to observe EhV-201 attachment and genome delivery (Fig. 2, S11, Movie S1). To facilitate the preparation of samples for electron-microscopy studies, we utilized *E. huxleyi* strain CCMP 2090, which does not produce coccoliths (Fig. 2, 3, S12) (*36*). In most cases, EhV-201 particles attached to *E. huxleyi* cells were oriented with one of their fivefold vertices towards the cell surface (Fig. 2BC, S11). We never observed capsids of EhV-201 entering cells. Some of the EhV-201 particles attached to the *E. huxleyi* surface contained emptied and partly collapsed inner membranes (Fig. 2C, S11E), indicating that they released their genomes by fusion of the inner virus membrane with the plasma membrane. This type of genome delivery requires the opening of the outer membrane and capsid of EhV-201. We hypothesize that the binding of the central and peripheral vertex proteins or of the putative filament to the host cell may trigger the conformational changes required for the outer membrane and capsid opening.

**Fig. 2.**
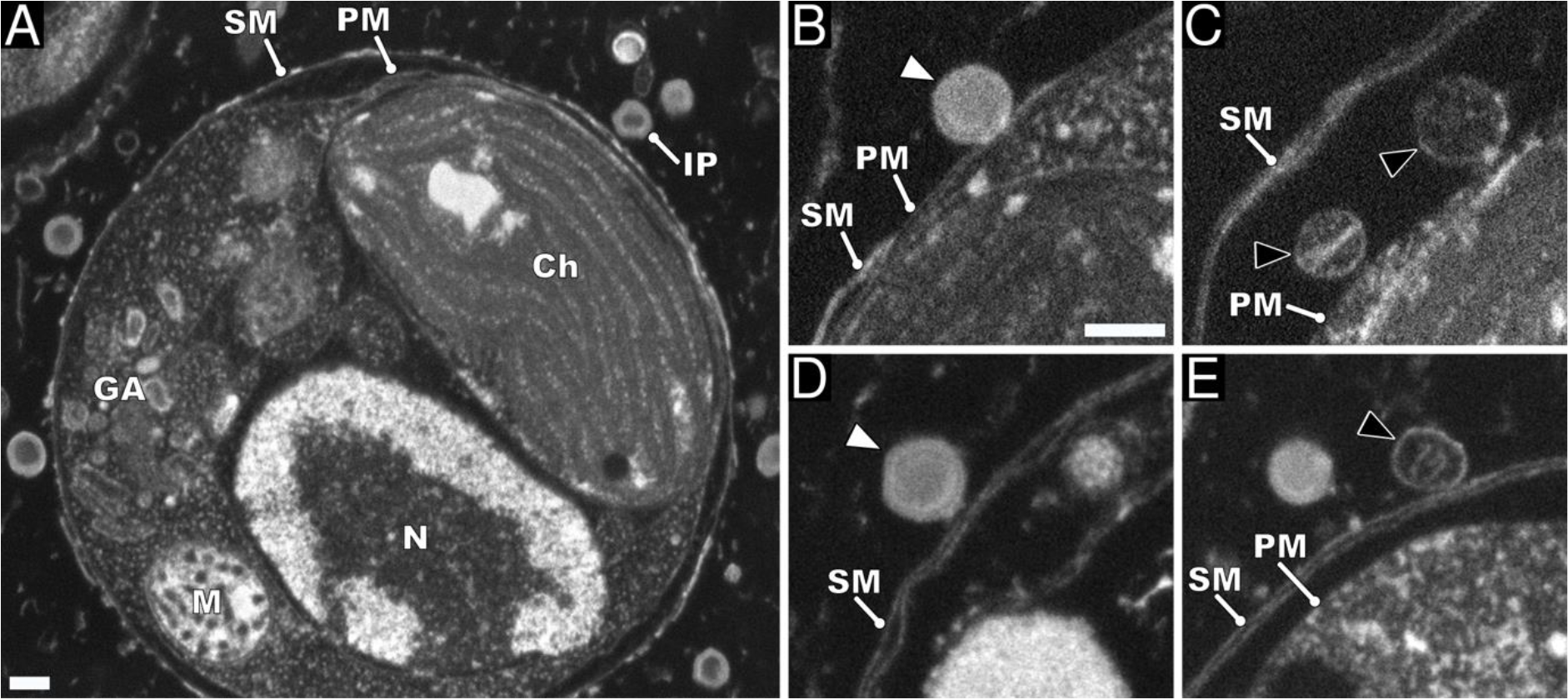
Attachment and genome delivery of EhV-201. **(A)** Scanning electron micrograph of high-pressure vitrified and resin-embedded *E. huxleyi* cell infected by EhV-201 at MOI = 10, 30 minutes post-infection. IP infecting particle, Ch chloroplast, GA Golgi apparatus, M mitochondrion, N nucleus, SM surface membrane, and PM plasma membrane. Scale bar 200 nm. **(B)** Genome-containing EhV-201 particle (white arrowhead) attached to plasma membrane of cell. Scale bar 200 nm. **(C)** Empty capsids (black arrowheads with white outlines) are attached to plasma membrane of cell. **(D)** Genome-containing EhV-201 particle attached to surface membrane of *E. huxleyi* cell. **(E)** EhV-201 particle that abortively released its genome after binding to surface membrane (black arrowhead with white outline).

**Fig. 3.**
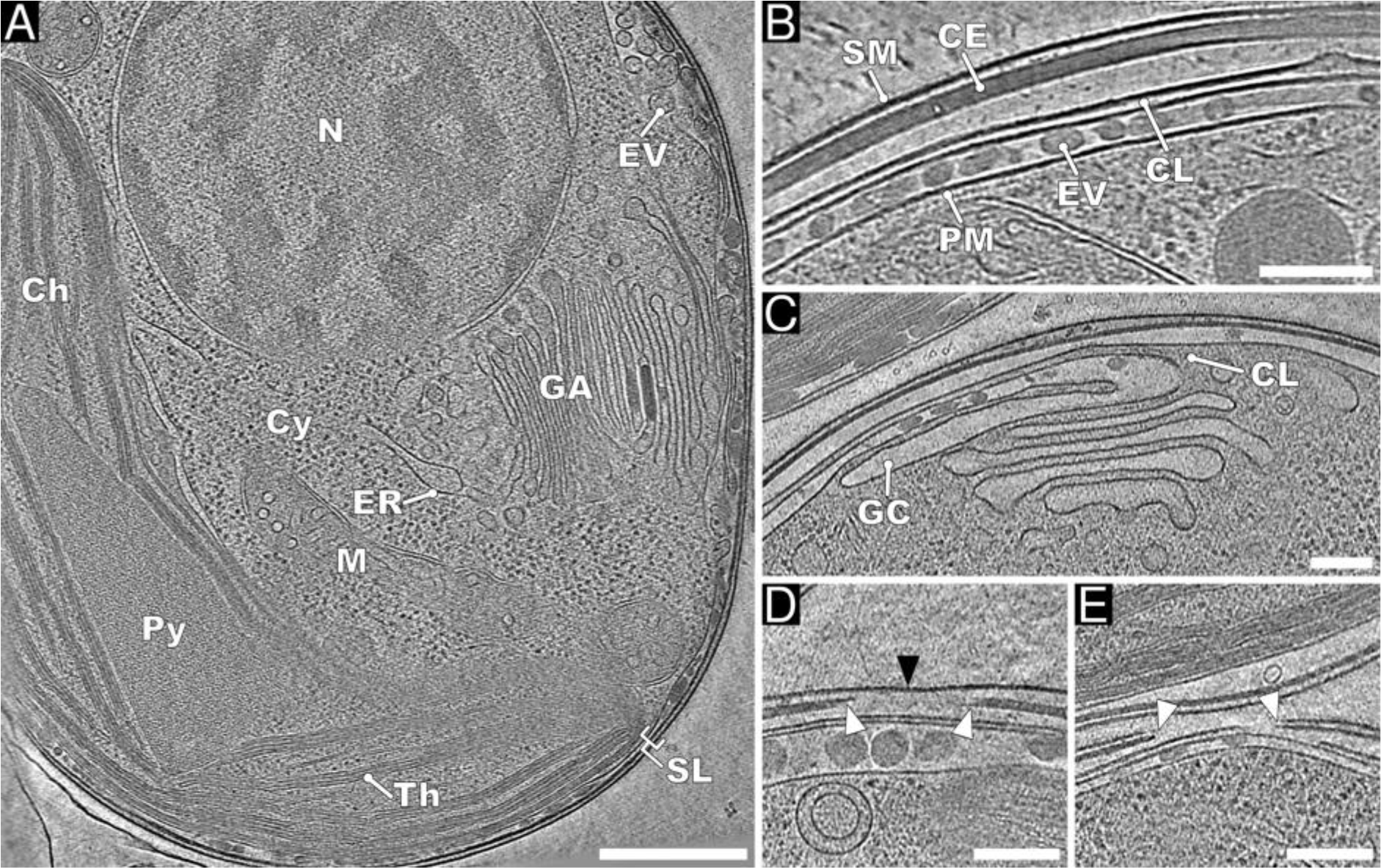
Native structure of *E. huxleyi* cell. **(A)** Projection image of 30-nm-thick section of cryo-tomogram of non-infected *E. huxleyi* cell from non-calcifying strain CCMP 2090. Scale bar 500 nm. **(B-E)** Details of organization of protective layers at *E. huxleyi* surface. N nucleus, Cy cytoplasm, CH chloroplast, Th thylakoid stacks, Py pyrenoid, ER endoplasmic reticulum, GA Golgi apparatus, M mitochondrion, and SL surface layers. Scale bars 200 nm. **(B)** Detail of continuous cell surface layers. EV extracellular vesicle between the PM plasma membrane and CL cytoplasmic leaflet, CE cell envelope, and SM surface membrane. **(C)** Cytoplasmic leaflet probably originates from fusion of GC Golgi apparatus cisternae with plasma membrane. **(D)** Opening in cell envelope, indicated by white arrowheads, is covered with a surface membrane (black arrowhead). **(E)** Opening in both the surface membrane and cell envelope makes plasma membrane accessible for virus infection.

### Cell surface layers protect *E. huxleyi* from EhV-201 infection

Most of the surface of *E. huxleyi* is covered with coccoliths, which, nevertheless, were shown to provide only limited protection against EhV infection (*37*). We used focused-ion-beam milling and cryo-electron tomography to show that EhV-201 virions can diffuse through the spaces in the coccolith structure (Fig. S13). Except for the missing coccoliths, CCMP 2090 cells are covered with the same surface layers that are found in wild-type cells: the surface membrane, cell envelope formed of polysaccharides, and cytoplasmic leaflets – large flat folds of a plasma membrane that wrap around the cell surface (Fig. 3, S13, Movie S2) (*38*). Cell envelope thickness ranges among the cells from 22 to 62 nm (mean 32 nm, SD = 13, N = 10), whereas the cell envelope of one cell is uniform in thickness (SD < 10%). We observed several EhV-201 particles that attached to or fused their inner viral membranes with the cell surface membrane, which resulted in the abortive release of the virus genomes into the extracellular space (Fig. 2D, E). This finding suggests that the surface membrane protects *E. huxleyi* cells from EhV infection. Furthermore, the cell envelope is impenetrable to virus particles. The cell envelope covered the surface of 93% (*n =* 43) of non-infected cells imaged using cryo-electron microscopy (Fig. 3); however, this was not resolved in the electron micrograph of resin-embedded cells (Fig. 2, S14). We speculate that the cell polysaccharide envelope could not be detected because it was not stained by the osmium tetraoxide and uranyl acetate used for sample contrasting (*39, 40*), or it could have been dissolved by the sample fixation procedure (*41*). The surface membrane and cell envelope contain openings that range in size from a few hundred nanometers in diameter to half of the cell surface (Fig. 3C, D). It is likely that the exposed areas of the plasma membrane render *E. huxleyi* cells sensitive to infection. The differences in the extent of cell coverage and, thus, in the protection provided by the surface membrane and envelope to individual *E. huxleyi* cells, make the EhV – *E. huxleyi* interaction complex at the population level. EhV-201 infection at a multiplicity of infection (MOI) of 10 did not clear the affected *E. huxleyi* culture after one virus replication cycle (Fig. S15). Only 1.4% (N *=* 211) and 19.4% (N *=* 227) of *E. huxleyi* cells had EhV-201 particles attached to their surface when infected with MOIs of 10 and 100, respectively (Fig. S16). Therefore, at an MOI of 100, only 0.2% of the virions capable of initiating an infection actually attached to the *E. huxleyi* surface. Our results agree with previous observations demonstrating that at most 25% of *E. huxleyi* cells from a population display infection symptoms at a given time (*42–45*).

### EhV-201 genomes replicate in the cell nucleus

After delivery into the cytoplasm, the genomes released from EhV-201 particles cannot be identified in images obtained using either serial block-face scanning electron microscopy or in cryo-tomograms of infected *E. huxleyi* cells. However, EhV-201 infection induces changes in the internal organization of the infected cells that enable indirect identification of the virus replication sites. Some NCLDVs, including Mimivirus and poxviruses, replicate their genomes in dense cytoplasmic replication factories (*46, 47*). In contrast, the remaining NCLDVs replicate in the cell nucleus, and their virus factories, which serve only for virion assembly, do not contain dense areas. Cryo-tomograms of EhV-201-infected *E. huxleyi* cells do not contain dense cytoplasmic virus factories, which corroborates previous evidence that the cocolithoviruses replicate in cell nuclei (Fig. 4) (*21, 22*). Whereas the nuclei of non-infected *E. huxleyi* cells contain distinct regions of heterochromatin and euchromatin and are enveloped by two membranes (Fig. 3A), this native nucleus morphology has never been observed in infected algae (N = 35) (Fig. 4A). The nucleus of an infected cell is characterized by a uniform distribution of its content and the absence of the outer nuclear membrane (Fig. 4A). Changes in the nuclear structure were reported previously for infections by NCLDVs that replicate in cell nuclei (*48, 49*).

**Fig. 4.**
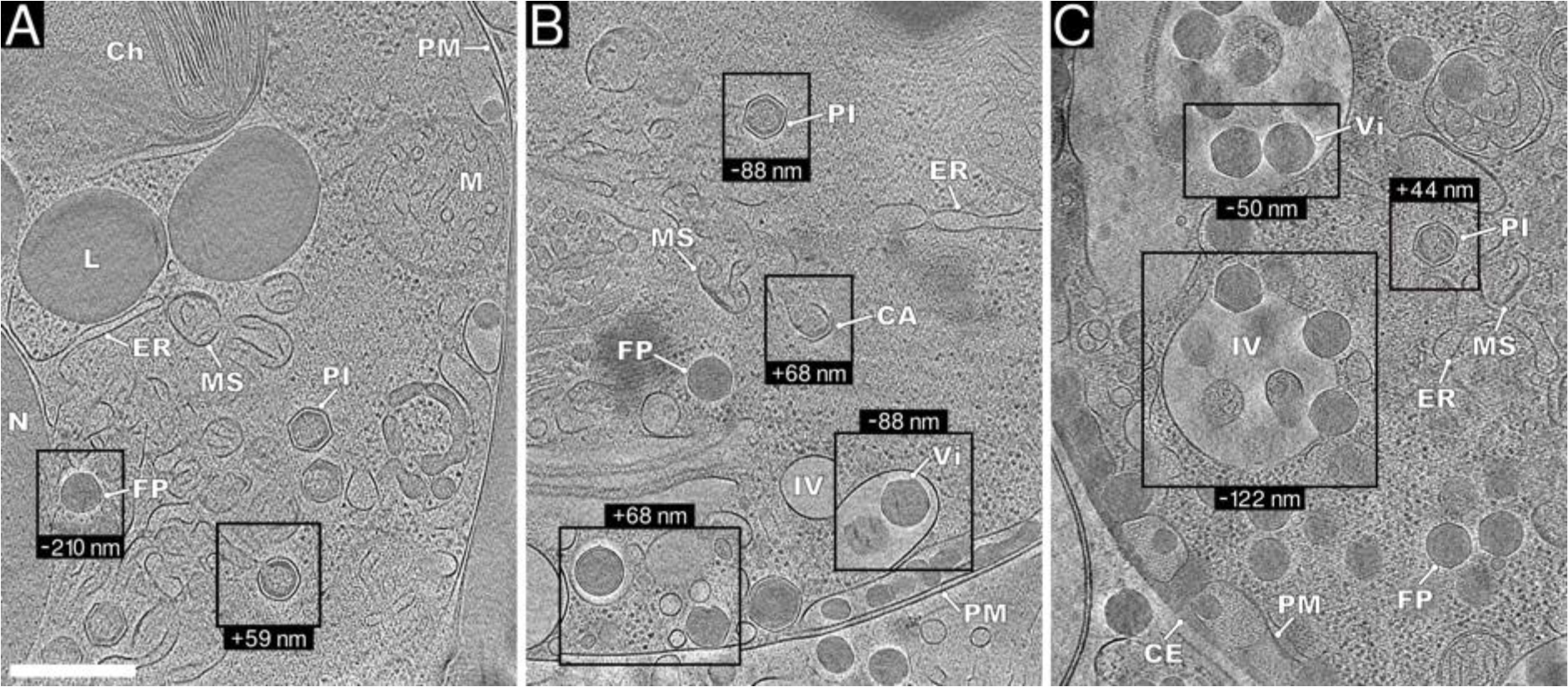
EhV-201 factories in *E. huxleyi* cells. **(A, B)** Virus factories in cells that lost surface protective layers could continuously release virions by exocytosis (80% of cells). **(C)** Virus factory with accumulated virions in cell with intact surface layers (20% of cells). The panels show projection images of 30-nm-thick tomogram sections of infected *E. huxleyi* cells. N nucleus, Ch chloroplast, M mitochondrion, ER endoplasmic reticulum, L lipid droplet, PM plasma membrane, and CE cell envelope. Components of EhV-201 factories: MS membrane segments, CA capsid assembly intermediate, PI genome packaging intermediate, FP full particle, Vi virion, and IV internal vesicle. Scale bar 500 nm.

### EhV-201 infection induces disruption of endoplasmic reticulum and outer nuclear membrane

The assembly of EhV-201 particles occurs in viral factories that occupy a segment of the cell cytoplasm between the nucleus and plasma membrane that is devoid of normal cellular organelles (Fig. 4). EhV-201-infected *E. huxleyi* cells do not contain the characteristic extensions of the endoplasmic reticulum, and their outer nuclear membranes are also partially or completely disrupted (Fig. 4). We speculate that EhV-201 infection induces the disintegration of the endoplasmic reticulum and outer nuclear membranes into segments that are the most abundant components of the viral factories (Fig. 4). The edges of the membrane segments are thermodynamically unfavorable structures that have to be stabilized by special proteins and lipids, the synthesis of which is probably ensured by EhV-201 (*17, 48, 50–52*). Molecular details of the mechanism stabilizing the membrane segments are as-yet unknown.

### EhV-201 capsid assembly and genome packaging

EhV-201 virion assembly initiates at a surface of a membrane segment located in the virus factory (Fig. 5). The early assembly intermediate consists of a membrane segment lined on one side by a featureless electron-dense layer (Fig. 5A, E), which has a similar appearance to that of incompletely packaged genomes inside assembling particles, and may therefore correspond to the initial stages of genome condensation (Fig. 5G). Alternatively, the electron-dense layer could represent scaffold proteins mediating the bending of the membrane segment. The face of the membrane segment opposite to that associated with the electron-dense layer serves as a nucleation site for capsid assembly (Fig. 5A, F). The forming capsids have straight edges and angular vertices, which indicate that they assemble according to the rules of quasi-icosahedral symmetry (Fig. 5B, G, H). An assembling capsid gradually encloses the membrane segment it is associated with and, in the process, induces its bending into a membrane sack (Fig. 5B, G). As the capsid assembly progresses, the virus DNA is packaged through an aperture in the forming capsid and the underlying membrane (Fig. 5B, H). When the assembly of the capsid nears its completion, the diameter of the DNA-packaging aperture in the capsid and membrane sack decreases to 15 - 40 nm (mean = 28; SD = 8; N = 6) (Fig. 5H). The capsid of EhV-201 can have several openings; however, we observed at most one aperture in the inner capsid membrane that served for genome packaging (Fig. 5G, H). During packaging, the EhV-201 genome forms condensed clusters inside the membrane sack (Fig. B, C, G, H). The mechanism of genome packaging of EhV-201, and probably also of other NCLDVs, is distinct from that of tailed bacteriophages and herpesviruses, in which the double-stranded DNA is pumped into the pre-formed capsid through a protein portal complex (*53, 54*).

**Fig. 5.**
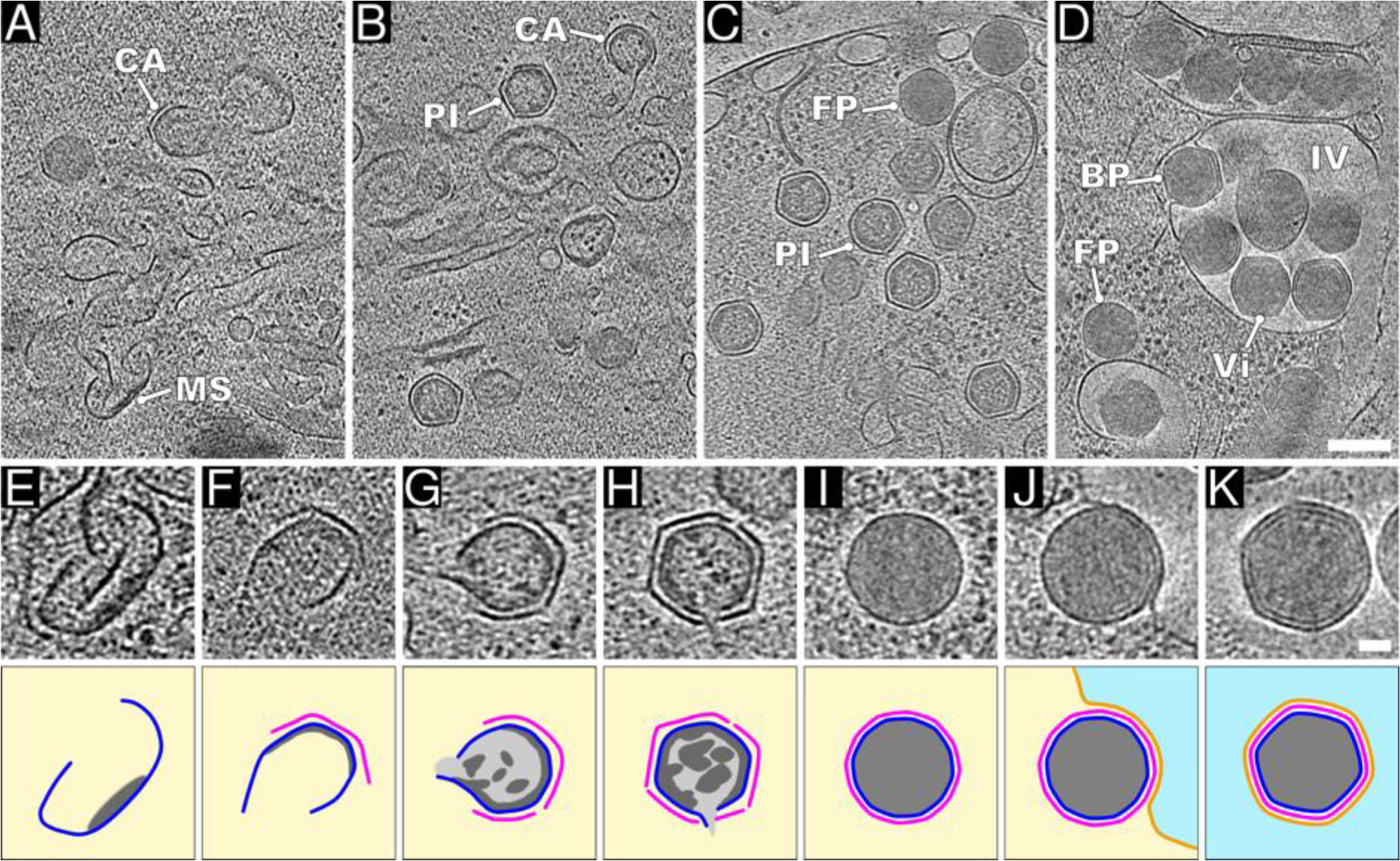
EhV-201 virion assembly pathway. **(A-D)** Virus factories contain assembly intermediates of EhV-201 virions. The panels show projection images of 30-nm-thick cryo-tomogram sections of infected *E. huxleyi* cells. (A) Degradation of host endoplasmic reticulum into MS membrane segments and CA capsid assembly intermediates. (B) Capsid assembly and genome packaging into PI packaging intermediate. (C) Conformational change of capsid and acquisition of uniform genome distribution occurs upon completion of genome packaging and formation of FP full particles. (D) Formation of Vi virions by budding of full particles into IV internal vesicles. Scale bar 200 nm. **(E-K)** Sections of cryo-tomogram reconstructions of assembly stages of EhV-201 particles with corresponding schematic drawings underneath. **(E)** Membrane segment (blue) with associated density that may correspond to condensing virus genome (dark grey). The yellow background indicates cytoplasm. **(F)** Capsid assembly is initiated at outer surface of membrane segment (magenta). **(G)** Early genome packaging intermediate with incomplete capsid and membrane sack. The genome has non-homogeneous distribution and is indicated by light and dark grey. **(H)** Genome packaging intermediate with inner membrane sack containing a single opening for DNA entry. Two openings in the capsid can be distinguished. The capsid has straight edges and angular vertices, indicating icosahedral symmetry. **(I)** A particle containing a complete genome. Due to conformational change, the capsid has become more oblique than that of the genome packaging intermediate, and the genome is homogeneously distributed. **(J)** Budding of full particles into an intracellular vesicle. The membrane of the vesicle is indicated by an orange line, and its interior by light blue. **(K)** Virion with complete outer membrane inside an intracellular vesicle. Scale bar 50 nm.

### EhV-201 particles acquire the outer membrane by budding into intracellular vesicles

The EhV-201 particles in the genome-filling stage are characterized by (i) angular capsids with a diameter of 193 nm (SD = 4), (ii) inner membranes separated from capsids, and (iii) clusters of packaged DNA (Fig. 5B, C, H, S17). When the genome packaging completes, the capsid becomes more spherical with a diameter of 190 nm (SD = 2) (Fig. S17), the inner membrane sack adheres to the capsid, and the genome distribution changes to homogeneous (Fig. 5C, I). Whereas the genome packaging intermediates exhibit no affinity for membranes, the genome-filled round particles bud into intracellular vesicles (Fig. 5D, J). Therefore, we speculate that the change in the capsid shape is connected to conformational changes in the major capsid proteins, which expose their amphipathic helices α3 and α4 from the DE-loop of the J1 domain at the particle surface and thus enable binding to membranes. This conformational change may also enable the binding of the capsid to EhV-201 transmembrane proteins, which must be present in the vesicles that the particles bud into. The budding of EhV-201 into vesicles is probably driven by the high-avidity interactions between the capsid and the vesicle membrane. The virions that completed budding and acquired an outer membrane have a diameter of 210 nm (SD = 4) (Fig. 5D, K, S17). Previous observations of EhV-201 and EhV-86 inside vacuoles support our hypothesis of EhV-201 budding into intracellular vesicles (*55, 56*). The budding process produces mature virions that need to be released from cells in order to initiate the next round of infection (Fig. 4, 5D, K).

### Exocytosis of EhV-201 virions

The formation of EhV-201 virions by budding into intracellular vesicles pre-disposes them to their release from the infected cells by exocytosis. However, the surface cell membrane and envelope cover 93% (*n =* 43) of native *E. huxleyi* cells and could block the diffusion of virus particles from the cell surface (Fig. 3). Efficient virion release is possible because EhV-201 infection induces the loss of surface membrane, envelope, and cytoplasmic leaflets from 80% (*n =* 35) of *E. huxleyi* cells (Fig. 4). In the sub-set of infected *E. huxleyi* cells that retained their surface membrane and envelope, EhV-201 virions accumulated beneath the protective layers (Fig. 4C, 6A, Movie S3). The budding into intracellular vesicles enables the continuous production and release of EhV-201 particles from the infected algae without the need for cell lysis. Previous studies by Mackinder *et al.* and Schatz *et al.* indicated that EhVs are released from cells by budding into the plasma membrane, based on electron microscopy images of thin sections of EhV-infected *E. huxleyi* cells (*21, 22*). However, the technique did not enable imaging of the cell envelope, and the sample preparation induced a shrinkage of cellular structures. It is therefore possible that the images presented by Mackinder *et al.* and Schatz *et al.* did not show virus budding, but instead corresponded to exocytosed particles trapped under the cell envelope, which may be more common in *E. huxleyi* cells covered by coccoliths (*21, 22*). Maturation processes involving budding into internal vesicles have been described for other enveloped NCLDV families that infect animals and amoebas, including *Asfaviridae*, *Poxviridae*, and *Mimiviridae* (*48*).

**Fig. 6.**
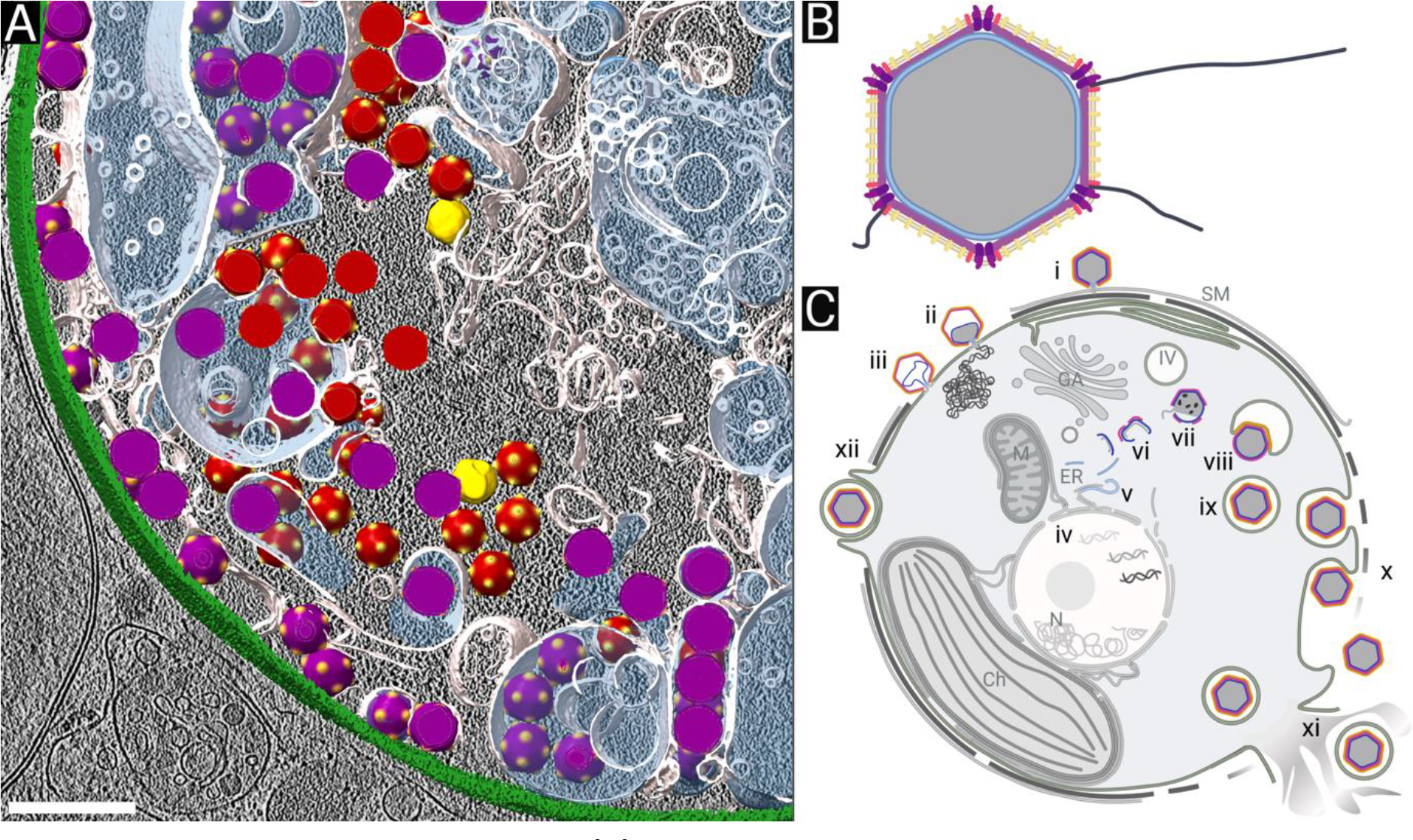
EhV-201 structure and replication. **(A)** Three-dimensional surface representation of segmented tomogram of an EhV-201-infected cell. The cell envelope is shown in green, cellular membranes in white, the content of intracellular vesicles is highlighted with semi-transparent blue, virions in red, full particles in orange, and assembly intermediates in yellow. Scale bar 500 nm. **(B)** Scheme of EhV-201 virion structure. The genome is shown in grey, the inner membrane in blue, the capsid in magenta, the outer membrane in orange, inner vertex proteins in purple, outer vertex proteins in red, ridge proteins in yellow, and fibers in black. **(C)** Infection cycle of EhV-201. Abortive infection: (i) Surface membrane and cell envelope protect *E. huxleyi* from EhV-201 infection. Productive infection: (ii) EhV-201 virion fuses its inner membrane with plasma membrane to deliver its genome into the cytoplasm. (iii) Empty capsid containing the collapsed inner membrane sack remains attached to the cell surface. (iv) EhV-201 genome probably replicates in the cell nucleus. (v) EhV-201 infection induces segmentation of the endoplasmic reticulum and outer nuclear membrane to form a virus factory. (vi) Genome packaging and capsid assembly initiate on opposite surfaces of membrane segments. (vii) Genome is packaged into a particle through an aperture in the forming capsid. (viii) The completion of the genome packaging induces a conformational change in the capsid, which enables it to bud into intracellular vesicles. (ix) Virion inside an intracellular vesicle. (x) EhV-201 infection causes the loss of surface protective layers from *E. huxleyi* cells, which enables the continuous release of virions by exocytosis. (xi) The EhV-201 replication cycle is terminated by cell lysis, which results in the release of virions inside vesicles. (xii) Alternative infection pathway utilizing phagocytosis of EhV virions inside vesicles. Panels B and C were created using BioRender.com.

### Vesicle-embedded EhV-201 virions released by cell lysis may infect cells when phagocytosed

The EhV-201 infection of *E. huxleyi* is terminated by cell lysis (*10, 21*), which also causes the release of virions embedded in vesicles (Fig. S18). We never observed infection of *E. huxleyi* by vesicle-bound EhV-201 particles; nevertheless, it has been shown that *E. huxleyi* cells in the late stationary phase phagocytose particles with a diameter of up to 500 nm (*57*). Therefore, the vesicle-bound virions may be phagocytosed by the alga, which could result in infection, as has been suggested by Mackinder *et al.* (*21*). Alternatively, the vesicle-bound particles could become infectious after the disruption of the vesicle membrane.

### Conclusions – EhV-201 structure and infection cycle

The EhV-201 virion initiates infection by binding to a cellular membrane using a particle vertex (Fig. 6B, C). Our results indicate that an EhV-201 particle delivers its genome into the cytoplasm by fusing its inner membrane with the plasma membrane of a cell (Fig. 6C). Attachment to the plasma membrane and opening of the EhV-201 capsid is probably mediated by transmembrane proteins positioned around the fivefold vertex of the particle or by a filament protruding from the vertex (Fig. 6B). Most *E. huxleyi* cells at a given time are resistant to infection by EhVs, probably because of the protection provided by the surface membrane and cell envelope that restrict access of the virus particles to the plasma membrane (*42*). The absence of dense structures in the cytoplasm of EhV-201-infected cells indicates that the virus genome replicates in the cell nucleus (Fig. 6A, C, Movie S3). The new particles assemble in virus factories located in the cytoplasm (Fig. 6A, C). Capsid assembly is initiated at the surface of endoplasmic reticulum-derived membrane segments (Fig. 6A, C). The genomes are packaged into the forming capsids through large apertures in the capsid and underlying membrane. After completion of the genome packaging, the capsids change their conformation, which enables them to acquire an outer membrane by budding into intracellular vesicles (Fig. 6A, C). EhV-201 infection induces the loss of the surface membrane and cell envelope from *E. huxleyi* cells, which enables the continuous release of EhV-201 virions by exocytosis (Fig. 6C). The EhV-201 replication cycle is terminated by cell lysis, which results in the release of virions inside vesicles from the ruptured cells (Fig. 6C). The vesicle-embedded virions can initiate infection if phagocytosed by *E. huxleyi* or after disruption of the vesicle membrane (Fig. 6C). Our study opens up numerous questions related to the molecular mechanisms of specific steps of the EhV-201 replication cycle such as: (i) What are the molecular interactions that enable the attachment of EhV-201 to the cellular membranes? (ii) How does the EhV-201 particle open to enable genome delivery? (iii) How does EhV-201 infection induce the loss of surface membrane and cell envelope to facilitate continuous virion release?

Characterization of the EhV-201 replication cycle contributes to our understanding of the general replication strategies employed by NCLDVs and highlights similarities in the nature of genome delivery and particle assembly across different viral families within the NCLDV group. *E. huxleyi* is a globally abundant marine phytoplankton species, and viral infections impact its population dynamics. Understanding the infection cycle of EhV-201, including its attachment, replication, and release mechanisms, can help to explain the dynamics of viral infections in marine environments and their potential consequences for changing marine ecosystems.

## Materials and methods

### Maintenance of *E. huxleyi* culture

*E. huxleyi* strain CCMP 2090 was cultivated in F/2-Si medium (*58, 59*) prepared as follows: seawater from an active marine aquarium (Aqua Vala, Brno, Czech Republic) was aged in the dark at 15°C for two weeks, passed through a 0.22 µm filter (Techno plastic products (TPP), Trasadingen, Switzerland) and further processed by tangential flow filtration (30 kDa MWCO, PES membrane; Pellicon XL 50; Millipore, Merck, Darmstadt, Germany), autoclaved, and enriched with micronutrients (Table S2). *E. huxleyi* cultures were inoculated to a final cell density of 2×10^5^ cells ml^-1^ in 600 ml tissue culture flasks (Jet BioFil, Guangzhou, China) and incubated in temperature- and illumination-controlled chambers (Photon Systems Instruments, Drásov, Czech Republic) at 15 °C, 50 µmol photons m^-2^ s^-1^ light intensity from LEDs with spectral ratios: white 33.3%; red 33.3%; far-red 33.3%, and a 16 h light / 8 h dark cycling regime with constant shaking (100 RPM, orbital shaker; N-Biotek, Bucheon, Republic of Korea). The cell density was measured using an automated cell counter (TC-20, Bio-Rad Laboratories, Hercules, California, USA) or by manual counting using a Bürker chamber (depth 0.1 mm, Thermo Fisher Scientific, Waltham, Massachusetts, USA).

### EhV-201 production and infectivity assays

EhV-201 (*19*) (Genbank accession code JF974311) was propagated on *E. huxleyi* strain CCPM 2090. An exponentially growing algal culture of *E. huxleyi* CCMP 2090 at a cell density of 1×10^6^ cells ml^-1^ was infected with EhV-201 at an MOI of 0.01 and left until complete lysis (up to one week). Viral stock solutions were prepared from the lysed algal culture by filtration through 0.22 µm syringe filters (Corning, New York, USA). The number of infectious viral particles was determined using plaque assay: 18 ml of an exponentially growing algal culture at a cell density of 1×10^6^ cells ml^-1^ was mixed with 100 µl of 10x serial dilutions of virus inoculum, incubated for 30 minutes at room temperature, mixed with 3% w/v low melting point agarose (UltraPure, Invitrogen, Thermo Fisher Scientific) in F/2-Si medium to a final concentration of 0.3%, and poured into Petri dishes (diameter 100 mm, Merck, USA). The cultures on Petri dishes were incubated in a translucent plastic box under the same conditions as the liquid algal cultures (*60*). Plaques, cleared round areas within the algae layer, were counted seven days post-infection.

### EhV-201 purification and preparation for cryo-EM single particle analysis

Two liters of an exponentially growing *E. huxleyi* CCMP 2090 at a cell density of 1×10^6^ cells ml^-1^ were infected with EhV-201 at an MOI of 0.01 and left until complete lysis (up to one week). For all subsequent purification steps, the lysate was kept on ice or at 4 °C. The lysate was sequentially filtered through a tangential flow filtration cassette with 0.45 µm pore size (PVDF membrane; Pellicon XL 50, Millipore) and concentrated on a 1,000 kDa MWCO tangential flow filtration cassette (regenerated cellulose membrane; Pellicon XL 50, Millipore). The concentrate was further concentrated using a centrifugal ultrafiltration unit (100 kDa MWCO, regenerated cellulose; Amicon, Merck) to 200 µl at 600 × g. The resulting concentrate was applied to a 10 - 50% v/v iodixanol step gradient with 10% increments (OptiPrep; Sigma Aldrich, Merck) enriched with sea salts (Sigma Aldrich) to a final concentration of 600 mM to maintain the salinity of F/2-Si media. Gradients were centrifuged in an ultracentrifuge (Optima XPN-80, Beckman Coulter, Danaher Corporation, Washington D.C., USA) using an SW-41 Ti rotor (Beckman Coulter) at 50,000 × g and 10 °C for 60 minutes. The 30 - 40% interface band was extracted using a needle and syringe (B.Braun, Melsungen, Germany) and dialyzed twice 1:1000 against aqueous sea salt solution (40 g L^-1^ Sigma Aldrich) in dialysis sleeves (15 kDa MWCO, Roth, Karlsruhe Germany). The dialyzed virus particles were concentrated in centrifugal ultrafiltration units to 200 µl and mixed with Turbo Nuclease (Abnova, Taipei, Taiwan) at a final concentration of 25 U ml^-1^ to digest free nucleic acids released into solution. Sodium azide (Sigma Aldrich) was added to a final concentration of 100 µg ml^-1^ to prevent bacterial growth. The purified virus was applied onto glow-discharged electron microscopy grids covered with holey carbon (Quantifoil, SPT Labtech, Melbourn, UK), blotted, and plunge-frozen using a Vitrobot Mark IV (Thermo Fisher Scientific) (Table S3).

### Cryo-EM data collection and single particle analysis

Data for single-particle analysis were acquired using a Titan Krios G2 cryo-TEM equipped with a Falcon 3EC direct electron detector (Thermo Fisher Scientific, Waltham, Massachusetts, United States) operating at 300 kV and a magnification at specimen level corresponding to a pixel size of 2.27 Å (Table S1) controlled with the software EPU 1.8 (Thermo Fisher Scientific). Motion correction of the original movies was done using MotionCor2 (*61*), CTF estimation of the aligned micrographs was performed using gCTF 1.06 (*62*), and particles were automatically picked using SPHIRE-crYOLO 1.7.5 (*63*) trained on a manually picked small dataset. The particle images were extracted from twofold binned micrographs with a 128 px box size and subjected to two-dimensional reference-free classification with a 500 Å diameter mask using Relion 3.1 (*64*). Class averages corresponding to either the virion vertices or the rounded surface areas, respectively, were selected for further analysis. To visualize the virion surface layers, pixel intensities along lines perpendicular to the particles’ surface were measured in the two-dimensional class averages using ImageJ 1.44 (*65*). Angular virion vertices were selected by three-dimensional (3D) classification using an initial model generated by the stochastic gradient descent method as implemented in Relion 3.1 (*64*). Subsequent refinement was performed using local searches with 1.8° sampling rate around the refined coordinates from 3D classification and applying a mask covering the capsid and outer membrane surface layers (Fig. S19).

### EhV-201 production, purification, and preparation for cryo-ET

A viral lysate was prepared in the same manner as the virions used for single-particle reconstruction, except for the addition of ampicilin and streptomycin (P-Lab, Praha, Czech Republic) (final concentrations of 100 µg ml^-1^) during algae culture cultivation and EhV-201 infection. Because the growth of the contaminating bacteria was reduced by the antibiotics, the OptiPrep step gradient and subsequent purification steps were omitted. Gold fiducials (6 nm; BSA tracer; Aurion, Wageningen, Netherlands) were buffer exchanged into 40 g L^-1^ sea salts (Sigma Aldrich) using centrifugal ultrafiltration unit (100 kDa MWCO, regenerated cellulose, Amicon) at 14,000 × g to the original volume. The centrifugal ultrafiltration concentrate of EhV-201 particles was mixed in a 3:1 ratio with the gold fiducials in sea salts. The final sample was applied onto electron microscopy grids covered with a holey carbon layer (Quantifoil), blotted, and plunge-frozen using a Vitrobot Mark IV (Thermo Fisher Scientific) (Table S3).

### Cryo-ET tilt series data collection, reconstruction, and sub-tomogram averaging

Tilt series were collected using a Titan Krios G2 cryo-TEM (Thermo Fisher Scientific) equipped with a K3 direct electron detector and energy filter (Gatan, Ametek, Berwyn, Pennsylvania, USA) operating at 300 kV at a magnification corresponding to a pixel size of 2.08 Å at specimen level (Table S1). Data were acquired using SerialEM 4.0 (*66*) and the protocol by Turonova *et al.* (*67*), with the following modifications: dose symmetric scheme starting from 0° with a 3° increment up to the maximum tilts of +/− 48° (Table S1). Original movies were motion-corrected using Warp 1.0.9 (*68*) and tilt-series were aligned using the IMOD 4.10.45 package (*69*). Viral vertices were picked using template matching in emClarity 1.5.3 (*70*) with the virion vertex from single-particle reconstruction as a reference structure. Sub-tomograms were extracted using Warp (*68*) from twofold binned images with a box size of 128 px, and imported into Relion 4.0 (*71*). Extensive 3D classification was performed using the virion vertex from the single-particle reconstruction as the initial model, applying a 500 Å wide circular mask and an additional mask covering the capsid and outer membrane or only the capsid, respectively. Final 3D refinement with local searches with 1.8° rotational sampling rate around the refined orientations from the 3D classification step and applying the same mask (Fig. S20). To achieve overlap between the density maps of neighboring vertices, the sub-volumes were re-extracted using threefold binning with a box size of 208 px. The larger vertices were reconstructed using local searches around coordinates from 500 Å 3D refinement with 1.8° rotational sampling and applying a circular 1,200 Å diameter mask and mask covering the capsid and outer membrane.

### Determination of EhV-201 virion *T*-number and generation of composite virion map

The *T*-number of the EhV-201 capsid was calculated by placing three sub-tomogram reconstructions of the vertices, calculated using a 1,200 Å diameter mask, into the tomographic volume at the positions and orientations determined from 3D refinement, using in-house developed scripts. The *h* and *k* values were counted as the number of steps along capsomers required to connect the pentons of two neighboring, partially overlapping vertex volumes.

To generate the composite map of the EhV-201 virion with idealized icosahedral symmetry, the density map of the vertex was rotated and translated to maximize the cross-correlation among its overlapping copies related by a threefold symmetry axis from the set of icosahedral symmetry axes in standard orientation, using an in-house developed script. The idealized composite map of the EhV-201 virion was prepared by expanding the aligned vertex density map according to the icosahedral symmetry.

### Sample preparation for cryo-focused ion beam milling

An exponentially growing culture of *E. huxleyi* CCMP 2090 at a cell density of 1×10^6^ cells ml^-1^ was infected with EhV-201 at an MOI of 10, and the infection was allowed to progress for 48 h. The infected cells were pelleted at 2,000 g for 10 min and resuspended in fresh F/2-Si medium at a final cell density of 1×10^7^ cells ml^-1^. Cells were applied onto electron microscopy grids covered with a holey gold layer (UltrAuFoil, Quantifoil, Jena Bioscience). The grids were flash-frozen using a Vitrobot Mark IV (Thermo Fisher Scientific) with backside blotting, for which the front-size blotting paper was replaced with a hydrophobic filter paper prepared in-house by soaking it in candle wax (Table S3).

### Cryo-lamellae preparation, tomographic data acquisition, and analysis

The cryo-lamellae were produced by cryo-focused ion beam milling (cryo-FIBM) using a Versa-3D dual beam microscope equipped with a gallium ion source (Thermo Fisher Scientific) and cryo-transfer chamber (Quorum Technologies Ltd., Lewes, United Kingdom). The sample was sputtered with conductive inorganic platinum (30 s at 10 mA) to produce 5-8 nm thick layer, and coated with 200 nm thick protective organic platinum layer (methylcyclopentadienyl platinum precursor, 15 s, cryo-deposited by the gas injection system heated to 28°C). Lamellae ranging in width from 5-8 µm were produced from clusters of vitrified cells. The rough lamella shape was achieved by parallel milling patterns with Ga^2+^ ions at 30 kV and 0.5 nA with a stage tilt of 15° (ion beam at 8° angle relative to the grid). The subsequent thinning and polishing of lamellae to a final thickness of less than 300 nm was performed at 100 pA and 10 pA, respectively. Lamellae were transferred to a Titan Krios G2 cryo-TEM (Thermo Fisher Scientific) equipped with a K3 direct electron detector with an energy filter (Gatan). Tilt series were collected using SerialEM 4.0 (*66*) and a dose-symmetric scheme starting from a pre-tilt of 8° with a 3° increment, covering relative tilt angles from −45° to +45° at a constant defocus of −25 µm. Magnification and the resulting pixel size of each data collection varied from 7.4 – 13 Å.

Tomograms of cryo-lamellae were reconstructed using IMOD 4.10.45. The individual tilts were aligned using patch tracking and the final binned tomograms were reconstructed at pixel sizes of 26 – 30 Å (*69*). The diameters of the viral particles and assembly intermediates were measured by fitting a circle around the most distal points of the virion using ImageJ 1.44 (*65*).

### Block-face imaging of resin-embedded algae

An exponentially growing culture of *E. huxleyi* CCMP 2090 at a cell density of 1×10^6^ cells ml^-1^ was concentrated as described in the cryo-lamellae FIBM preparation section, and resuspended in the F/2-Si medium at a cell density of 1×10^9^ cells ml^-1^. Cells were mixed with EhV-201 concentrate prepared in the same manner as that for sub-tomogram averaging (without the addition of sodium azide) to obtain an MOI of 10. The mixture was incubated for 30 min at room temperature. Cryo-samples were prepared using high-pressure freezing in 200 µm carriers (cavity volume 0.6 mm^3^) treated with 0.2% w/v lecithin in chloroform (Sigma Aldrich) using an EM ICE device (Leica Microsystems GmbH, Wetzlar, Germany) at 2,010 bar. Cryo-preserved samples were freeze-substituted with 1% w/v osmium tetraoxide (Sigma-Aldrich, Cat.-No. 45345) in acetone (Penta, Chrudim, Czech Republic) using an EM AFS2 device (Leica Microsystems) using protocol: −90 °C for 16 h; −90 °C to −85 °C for 6 h; −85 °C to −60 °C for 6 h; −60 °C for 5 h; −60 °C to −30 °C for 6 h; −30 °C for 4 h; and −30 °C to +10 °C for 6 h. Sample was further post-stained with 1% uranyl acetate (Electron Microscopy Science, PE, USA) infiltrated with epoxy embedding medium (Sigma-Aldrich) at Epoxy:acetone ratio of 30:70 for 2h; 50:50 for 2 hours; 70:30 for 2 hours; and 4 times with a fresh Epoxy medium for 12 h. Polymerization was allowed to progress at 65 °C for 48h. Resin blocks were mounted on an aluminum SEM stub and sputter-coated with 5 nm platinum using a Quorum Q150T (Quorum Technologies Ltd.). The serial block milling and imaging were performed using a Helios Hydra 5 CX with Auto Slice & View v. 4.2 (Thermo Fisher Scientific). A platinum protective layer (methylcyclopentadienyl platinum precursor) was deposited by a gas injection system on top of the region of interest to a total thickness of 1 µm at 30 kV and 1 pA. Trenches were milled using a xenon plasma beam at 30kV and 15nA, resulting in a final block size of 45 x 45 x 50 µm. In order to reduce the curtaining artifact, oxygen plasma (30kV and 30pA) was used for serial milling. Slices of a final size of 45 µm x 50µm x 20nm were imaged using an ICD detector in immersion mode (2 kV and 200 pA, dwell time 10 µs). The resulting images had a resolution of 3072 x 2048 px with a pixel size of 1.69 nm. The final data cube was aligned, stitched without further post-processing, and analyzed using ImageJ 1.44 (*65*).

### Fluorescence microscopy

Exponentially growing cultures of *E. huxleyi* CCMP 2090 at a cell density of 1×10^6^ cells ml^-1^ were concentrated as for cryo-FIBM sample preparation and resuspended in F/2-Si medium at a cell density of 1×10^7^ cells ml^-1^. Concentrated EhV-201 virions, prepared as for sub-tomogram averaging (without the addition of sodium azide), were stained overnight with 4′,6-diamidino-2-phenylindole (DAPI, Roche, Basel, Switzerland) at a final concentration of 5 µg ml^-1^. Excess stain was removed by dialysis at a 1:1000 ratio against aqueous sea salt solution (40 g L^-1^, Sigma Aldrich) using dialysis sleeves (14 kDa MWCO, Roth) at 4°C in the dark. Algae were mixed with EhV-201 at MOIs of 10 and 100, respectively. The mixtures were incubated at room temperature in the dark for 30 min, stained with N-(3-triethylammonium propyl)-4-(4-(dibutyl amino) styryl) pyridinium dibromide (FM 1-43FX, Invitrogen, Cat.-No. F35355) at 5.6 µg ml^-1^. After 5 minutes of incubation, the mixture was fixed using glutaraldehyde (Penta, Chrudim, Czech Republic) at a final concentration of 0.2% v/v and immediately transferred onto a 10-well cell culture microscopy slide (CellView; Greiner Group AG, Kremsmünster, Austria) pre-treated with poly-lysine (Sigma Aldrich) dissolved in distilled water at a concentration of 0.01% w/v. After a 30 min settling time, the supernatant was removed, and the wells were covered with mounting media (ProLong glass antifade, Invitrogen).

Fluorescence images were recorded using an Elyra 7 super-resolution microscope operated in lattice structured illumination microscopy (SIM) mode and controlled using the ZEN black edition 3.0 system (ZEISS, Oberkochen, Germany). Volume data were acquired using an oil immersion objective (Plan-Apochromat 63x/1.4 Oil DIC) and detected with a pco.edge sCMOS camera using a frame size of 512 px (x,y) with 9-phase grating in leap mode using the following channel settings: DAPI: 405 nm laser 20% excitation, BP420-480 dichroic mirror and SBS BP 490-560 beam splitter, exposure time 50 ms; FM 1-43: 488 nm laser 1% excitation, 495-590 dichroic mirror and SBS BP 490-560 beam splitter, exposure time 30 ms. Acquired volume data were reconstructed with ZEN black edition 3.0 (ZEISS) using the 3D SIM^2^ method with drift correction measured using fluorescent beads.

The distance of the virus particle from the algal cell surface was determined by subtracting the radius of a circle fitted around the circumference of the (FM 1-43-stained) algal plasma membrane from the larger circle with the same origin crossing the viral DAPI-stained genome. The cut-off distance for attachment was set to 300 nm, i.e., 1.5 times the virus particle diameter.

### Reconstruction and segmentation of tomograms of infected *E. huxleyi* cells

Tilt-series of cryo-lamellae were reconstructed using the package IMOD 4.10.45 (*69*). Selected tomograms of infected and non-infected control cells were segmented using artificial intelligence-assisted segmentation as implemented in DragonFly 2022.2.0 (Object Research System, Montréal, Québec, Canada). The network was trained on a small portion of manually segmented tomogram volume and applied to the entire tomogram, followed by extensive manual pruning. The positions of EhV-201 virions and full particles were determined by template matching using emClarity 1.5.3.11 (*70*) and composite maps of EhV-201 virions with and without the outer membrane, respectively, were placed back into the tomographic volume using in-house developed scripts. Images and movies of the segmented tomograms were generated using ChimeraX 1.5 (*72*).

### Size comparison of distinct EhV-201 assembly intermediates

Statistical analyses were performed using the R v4.2.2 in RStudio environment v2022.07.2 and the following libraries: emmeans_1.8.1-1, car_3.1-0, carData_3.0-5, lme4_1.1-30, and Matrix_1.5-1 (*73–75*). Data visualization was performed using ggplot2_3.3.6 (*76*). The maximum-outer diameters of virus particles were measured in tomographic reconstructions of infected *E. huxleyi* cells: genome packaging intermediate (N = 25), full particle (N = 25), and virion (N = 25) in triplicates and averaged. The diameter of each particle was measured at different Z-heights to determine the maximum diameter. Particle diameter was treated as a continuous response variable, and assembly stage was treated as a fixed factor. The Dataset_ID (3 levels) of different sample preparations and data collections were treated as random factors because cell_ID (9 levels), an identifier of individual cells in the cryo-lamellae, did not contain all the assembly stages equally distributed. The diameter of particles was compared by a linear mixed model (LMM) with dataset_ID as a random factor. The H0 of equal particle diameter was rejected by analysis of variance (ANOVA Type-II) at p < 0.0001 (Df = 73; F = 229.6). Pairwise analysis of diameter between intermediate stages to determine the p-value was done by multiple analysis of means (MANOM).

### Effect of DAPI-staining on EhV-201 infectivity

The titer of a viral lysate was determined from the concentration of plaque-forming units using plaque assay (N = 3). The concentration of plaque-forming units was treated as a continuous response variable, and DAPI treatment as a predictor variable. The concentrations of plaque-forming units were compared using the Welch two-sample (two-tailed) t-test, and the H0 of an equal number of plaques was not rejected at p = 0.195 (Df = 4, t = 1.63).

## Acknowledgments

We gratefully acknowledge (i) the Cryo-electron Microscopy and Tomography Core Facility and Proteomics Core Facility of the Central European Institute of Technology (CEITEC), Masaryk University, supported by the Ministry of Education, Youth, and Sports of the Czech Republic (Grant LM2018127); (ii) the Cellular Imaging Core Facility supported by the Czech-BioImaging large RI project (LM2018129 funded by MEYS CR); and (iii) Plant Sciences Core Facility for their support with obtaining scientific data presented in this paper. We gratefully acknowledge support from the project National Institute of Virology and Bacteriology (Program EXCELES, ID Project No. LX22NPO5103) - Funded by the European Union - Next Generation EU. This work received funding from the Czech Science Foundation Grant GX 19-259882X to P.P. and from European Regional Development Fund-Project „MSCAfellow2@MUNI“ (No. CZ.02.2.69/0.0/0.0/18_070/0009846) to C.R.B. and Brno Ph.D. talent scholarship funded by Brno city municipality to M.H. We are thankful to Jakub Zak (ORCID 0000-0003-2845-8323) for his help with statistical analyses.

## Data and code availability

Cryo-EM maps and structure coordinates were deposited to the Electron Microscopy Data Bank (EMDB) and Protein Data Bank (PDB), respectively, with the following accession numbers: (i) EhV-201 virion vertex reconstructed using a single particle approach and a mask with the diameter of 50 nm – EMD-17650; (ii) EhV-201 virion vertex reconstructed using sub-tomogram averaging and a mask with the diameter of 50 nm – EMD-17649, and (iii) EhV-201 virion vertex reconstructed using sub-tomogram averaging and a mask with the diameter of 50 nm and a mask covering only the capsid layer – EMD-17648, the fitted structure of AlphaFold2 predicted EhV-201 major capsid protein and PBCV-1 penton protein – PDB 8PFM; (iv) EhV-201 virion vertex reconstructed by sub-tomogram averaging and a mask with the diameter of 120 nm - EMD-17651. All custom scripts used during tomographic reconstruction have been made available at https://github.com/fuzikt/tomostarpy.

## Author contributions

Major contributions to (i) the conception and design of the study M.H., C.R.B., D.C.S., and P.P.; (ii) the acquisition, analysis, and interpretation of the data M.H., C.R.B., T.F., P.K., R.H., J.N., M.C., F.F., W.H.W., D.C.S., and P.P.; and (iii) the writing of the manuscript M.H., C.R.B., and P.P. All authors commented on the manuscript.

## Supplementary figures

**Fig. S1.**
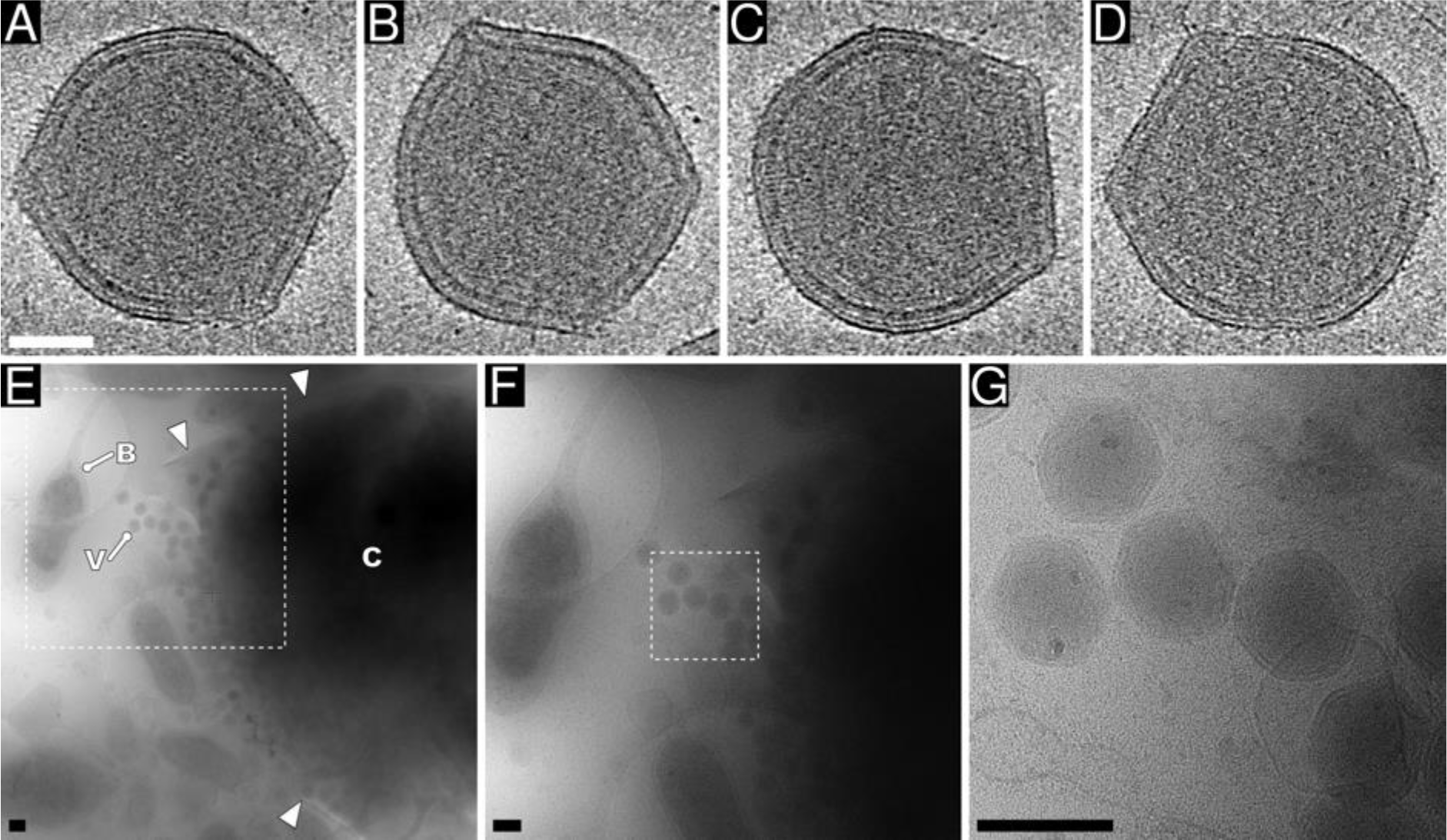
EhV-201 virions are pleomorphic. **(A-D)** Projection images of 8-nm-thick sections from cryo-tomograms of EhV-201 virions, which show regions with sharp edges and angular vertices, but also rounded parts. The particles differ from each other. Scale bar 50 nm. **(E-G)** Cryo-electron micrograph of EhV-201-infected *E. huxleyi* cell that lysed during the vitrification of the sample for cryo-EM. Scale bar 200 nm. (E) Overview of lysed cell. C remnants of *E. huxleyi* cell, B bacteria that grow in co-culture with *E. huxleyi*, V EhV-201 virion. White arrowheads indicate the edges of the ruptured plasma membrane. The dashed square indicates the position of the region shown at a higher magnification in panel (F). (F) Intermediate magnification of virions released from ruptured *E. huxleyi* cell. The dashed square indicates the position of the region shown at higher magnification in panel (G). (G) EhV-201 virions are deformed and differ structurally from each other immediately after release from lysed cell.

**Fig. S2.**
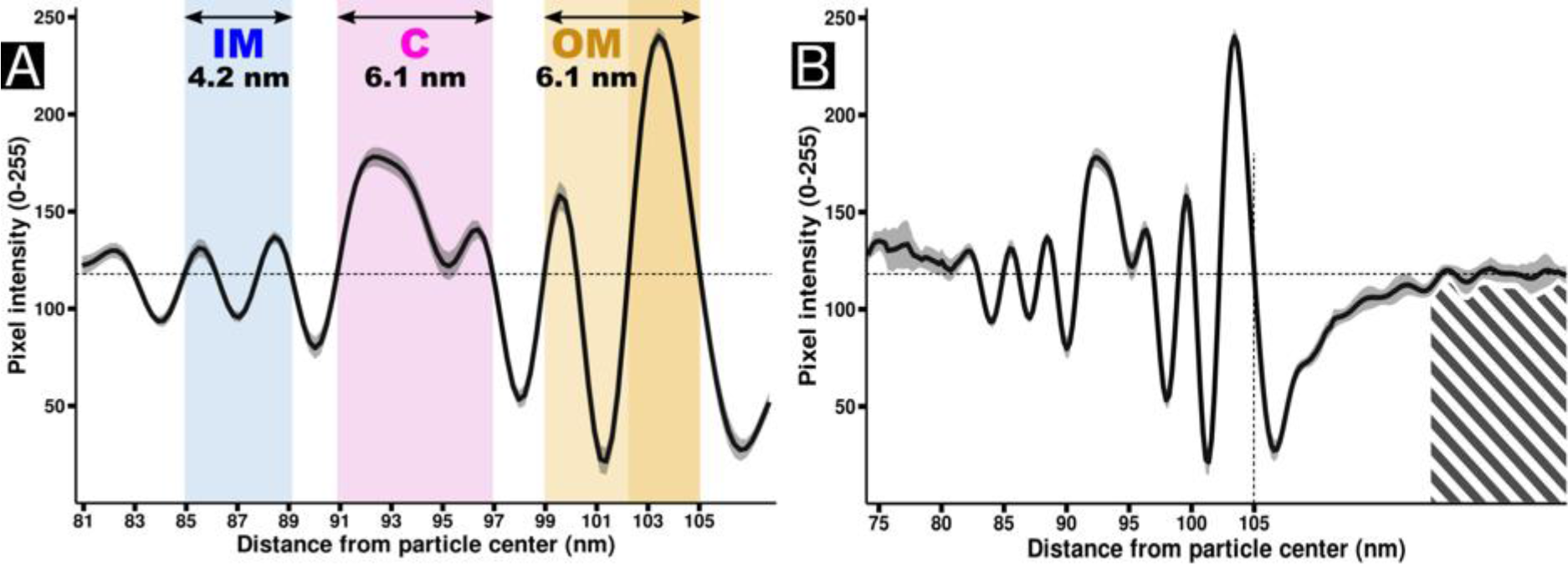
Surface layers of EhV-201 virion. **(A)** Plot of average pixel intensities measured along lines perpendicular to particle surface in reference-free two-dimensional class average of oblique segments of EhV-201 particle surface. Layers representing the inner membrane, capsid, and outer membrane are indicated by colored backgrounds (IM inner membrane is shown in blue, C capsid in magenta, and OM outer membrane in orange). Numbers indicate layer thickness. The outer leaflet of the outer membrane, indicated by dark orange, has a stronger density than the inner leaflet. The coloring scheme corresponds to that in Fig. 1B, C. The 95% confidence interval is indicated by grey shading. The average background intensity level is indicated by a horizontal dashed line. **(B)** Extended plot including region used for determination of average background value calculated from hatched area. N = 18.

**Fig. S3.**
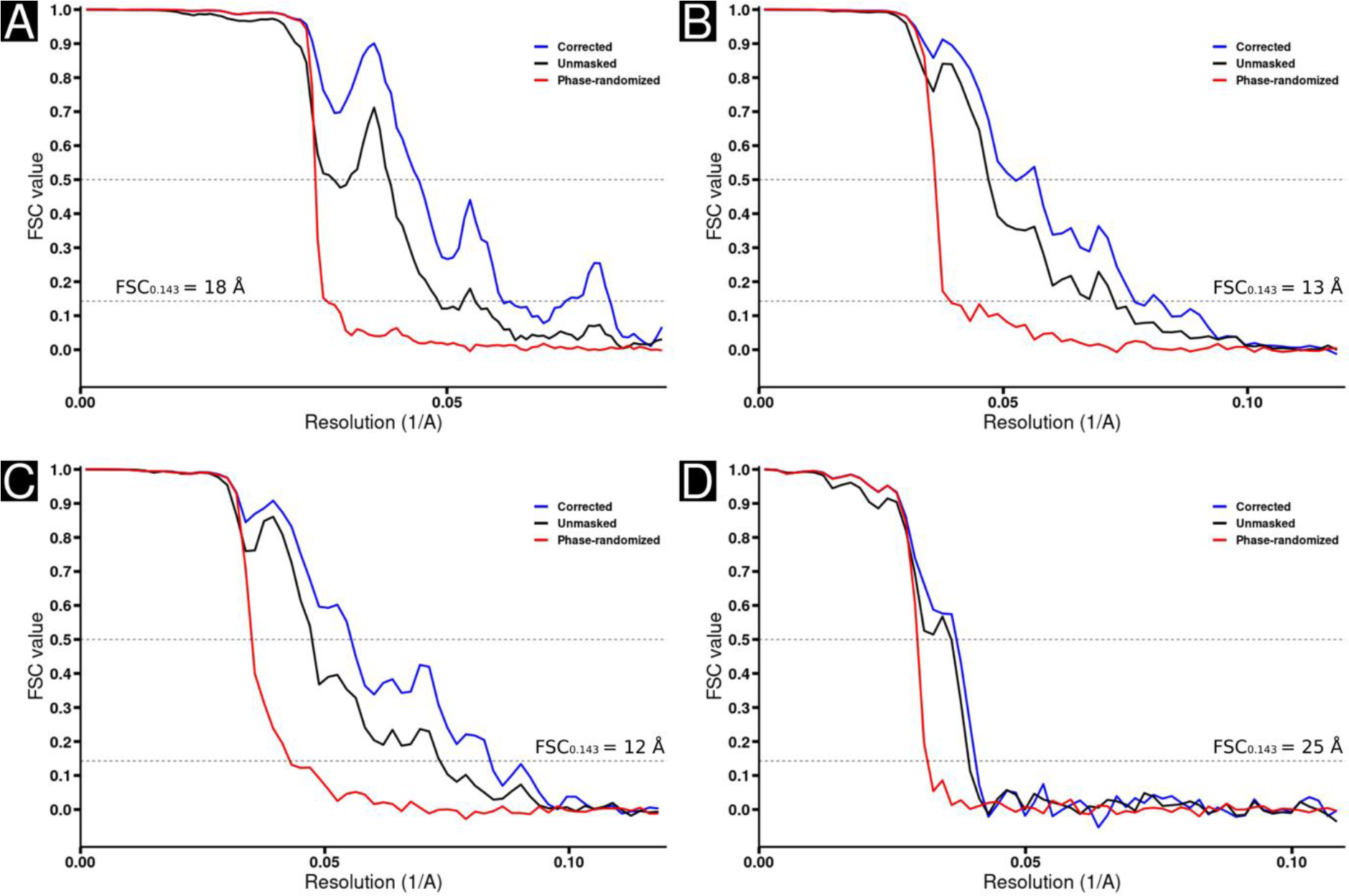
Plots of Fourier shell correlation (FSC) of reconstructions of independent halves of cryo-ET and cryo-EM datasets of EhV-201 virion vertices. **(A-B)** Sub-tomogram reconstructions of EhV-201 virion vertex with masks limiting the size of the reconstruction to 120 nm (A) and 50 nm (B). **(C)** Sub-tomogram reconstructions of EhV-201 virion vertex with mask limiting the size of the reconstruction to 50 nm and removing the outer and inner membrane. **(D)** Single-particle reconstruction of EhV-201 virion vertex with masks limiting the size of the reconstruction to 50 nm. Dashed lines indicate FSC values 0.5 and 0.143.

**Fig. S4.**
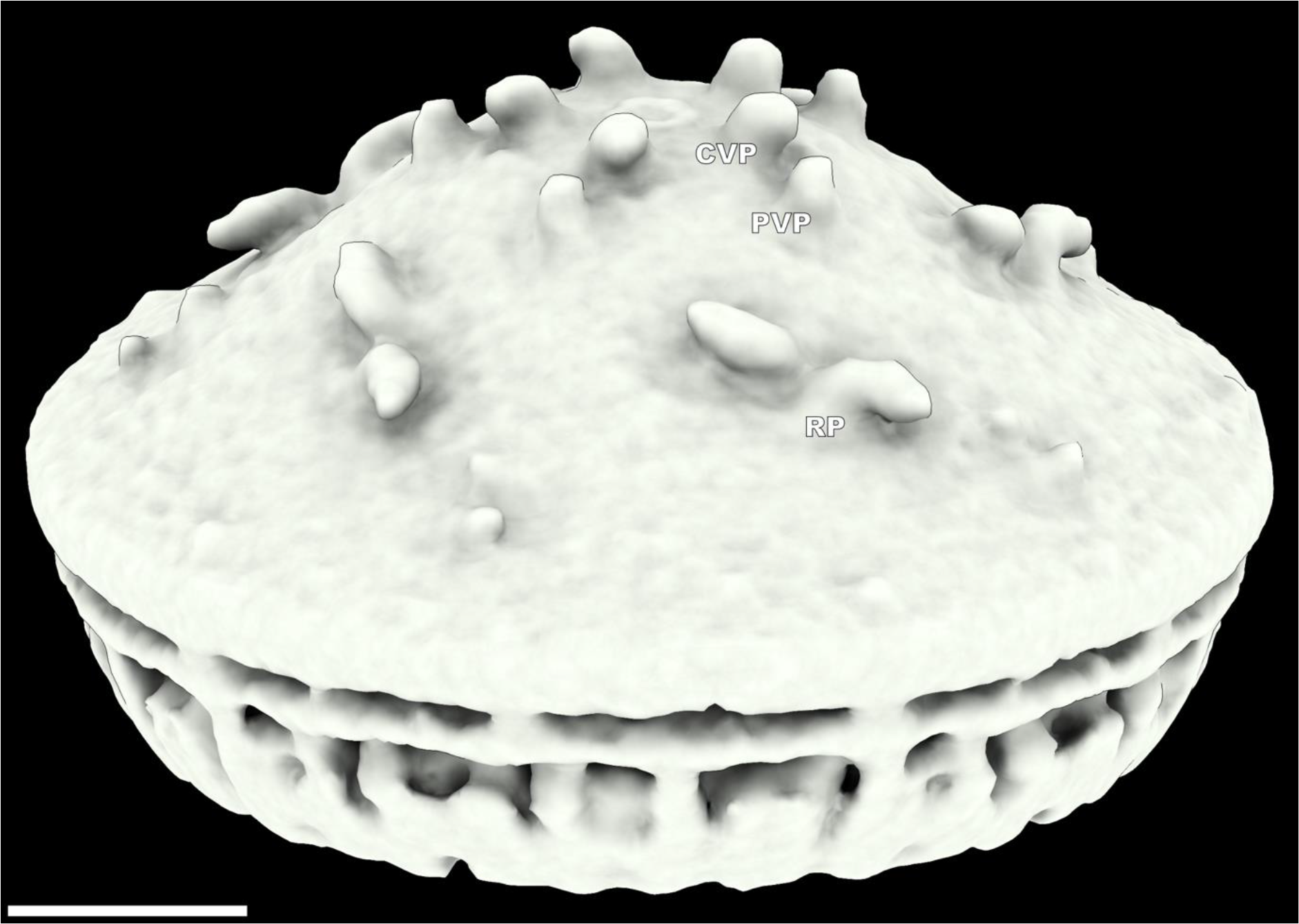
Outer membrane of EhV-201 is decorated with transmembrane proteins. Surface representation of sub-tomogram of EhV-201 vertex reconstructed using a mask with the diameter of 50 nm showing central vertex proteins (CVP), peripheral vertex proteins (PVP), and dimers of ridge proteins (RP). Scale bar 10 nm.

**Fig. S5.**
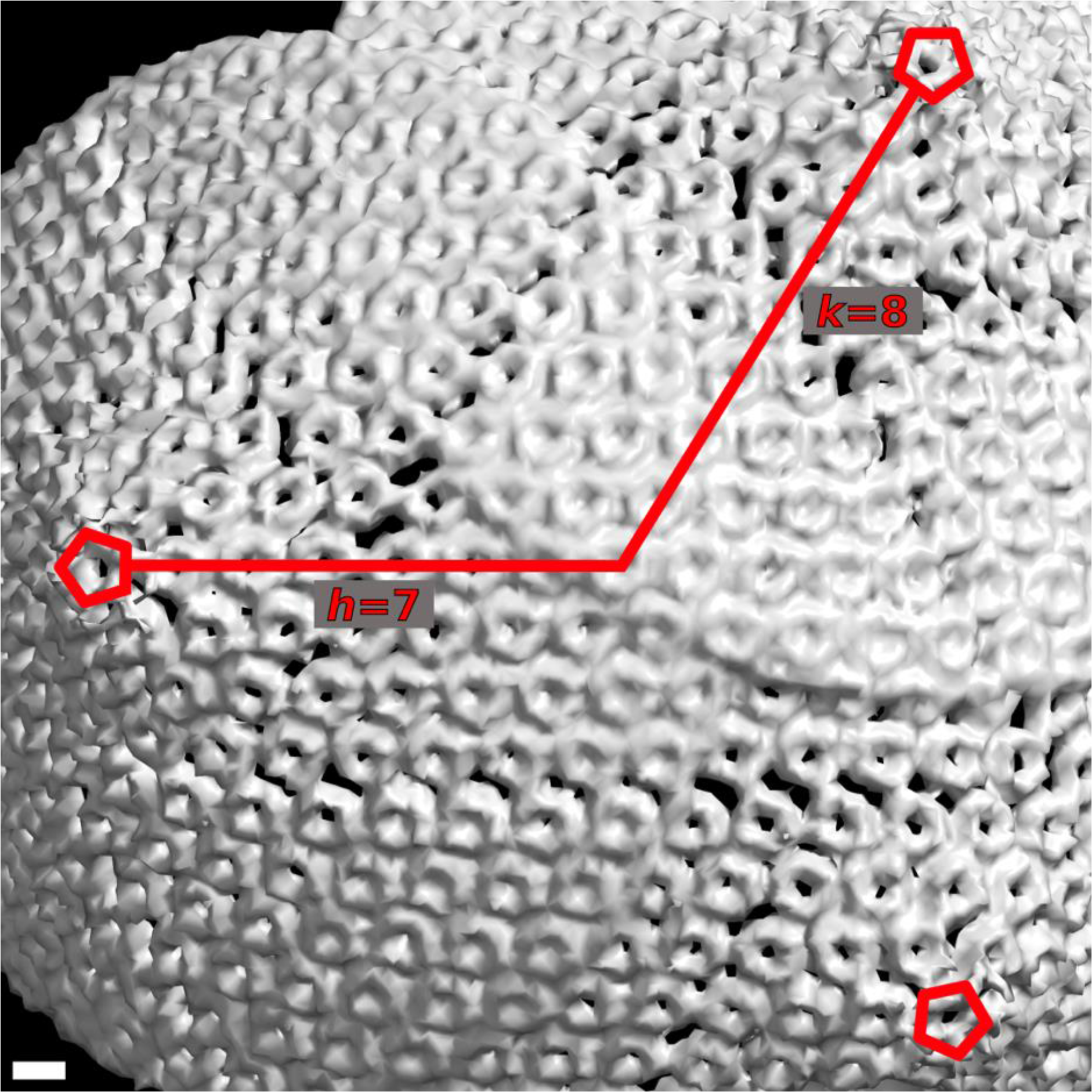
EhV-201 capsid is organized with *T* = 169 quasi-symmetry. Surface representation of three cryo-ET reconstructions of angular vertices (mask diameter 120 nm) placed back into tomogram of EhV-201 virion based on coordinates obtained by three-dimensional refinement. Positions of fivefold symmetry axes are indicated by red pentagons, *h* and *k* directions are indicated by red lines. Scale bar 5 nm.

**Fig. S6.**
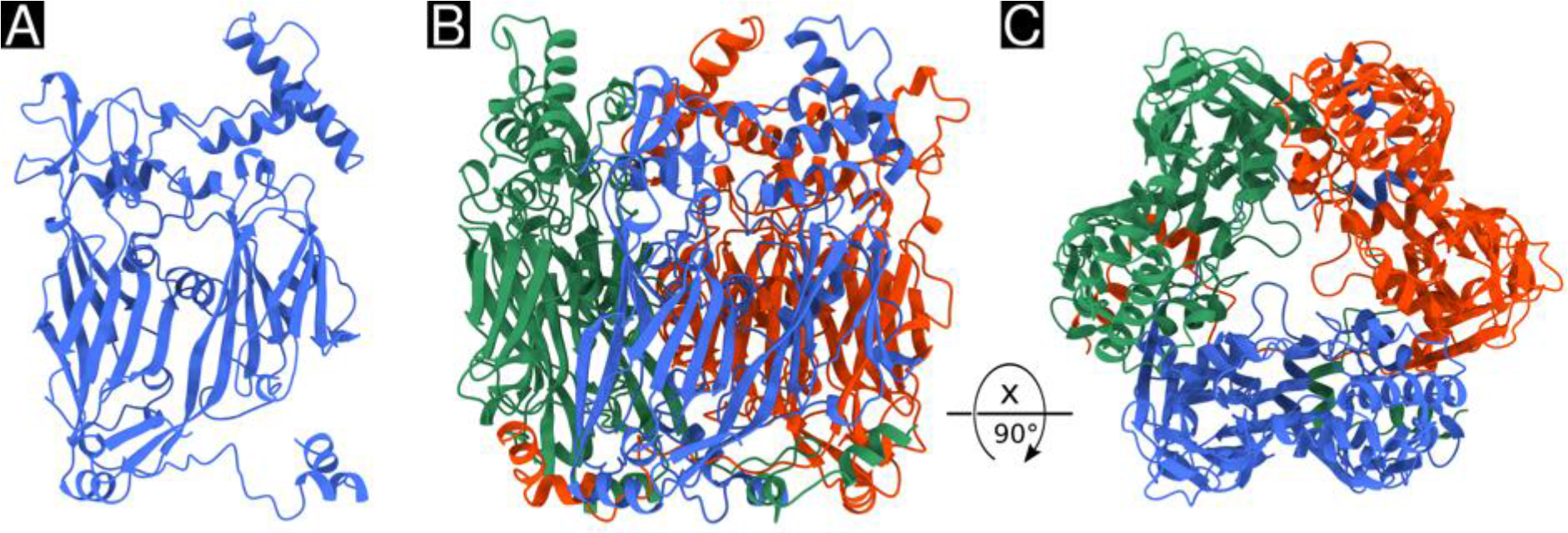
Structure of EhV-201 major capsid protein and capsomer. **(A)** Cartoon representation of AlphaFold2 (*27*) predicted structure of EhV-201 major capsid protein. J1 and J2 indicate the two jellyroll domains of the major capsid protein. **(B, C)** Side (B) and top (C) view of capsomer formed by three major capsid proteins shown in red, green, and blue.

**Fig. S7.**
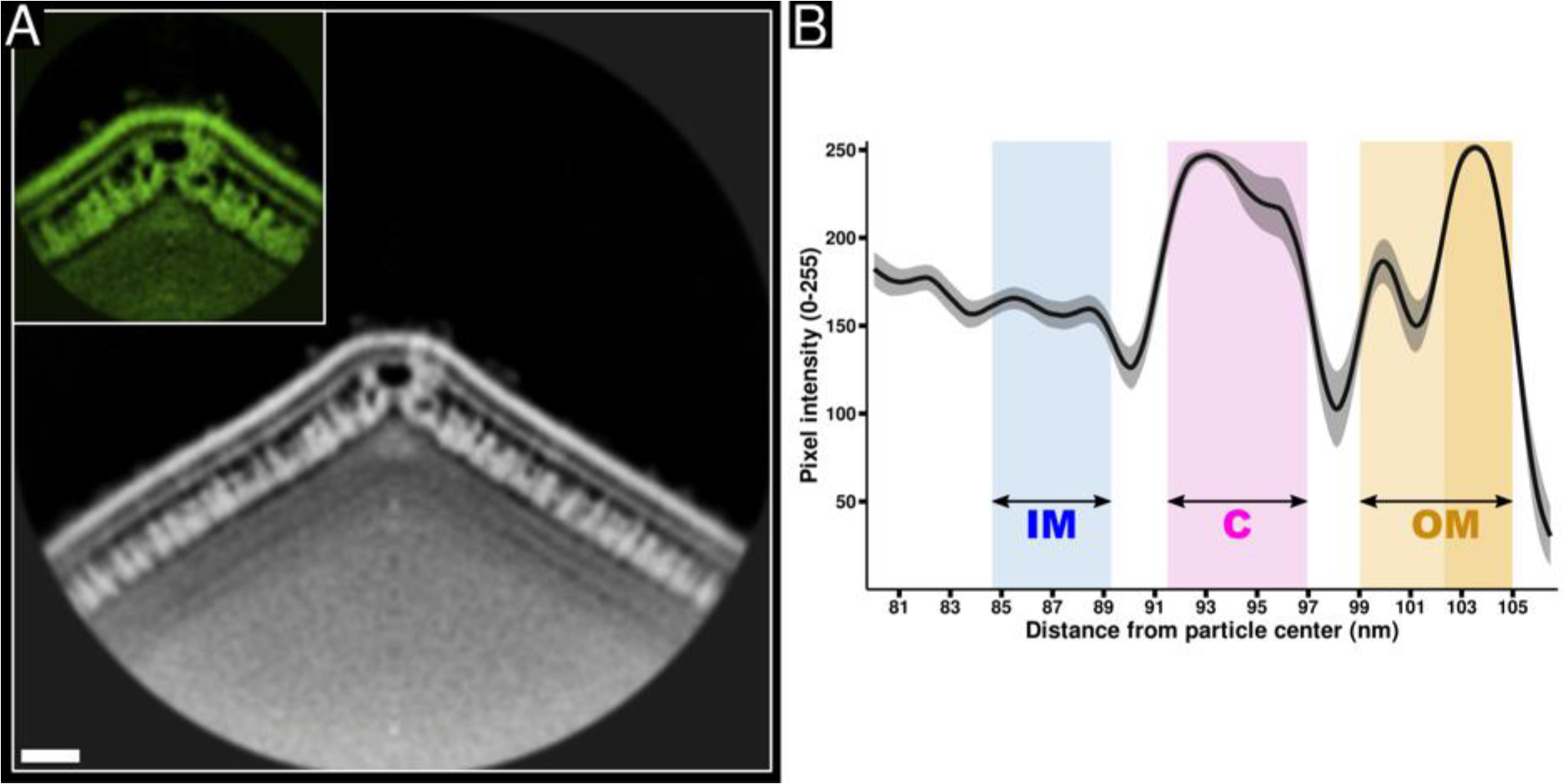
Inner membrane is less resolved than outer one in EhV-201 virion vertex sub-tomogram reconstruction. **(A)** Central sections of sub-tomogram reconstructions of vertices from EhV-201 virion with reconstruction diameter set to 120 nm (grey) and 50 nm (inset in green). No feature-based masks were applied in the reconstruction process. Scale bar 10 nm. **(C)** Plot of average voxel intensities measured along lines perpendicular to EhV-201 virion surface. Layers representing the inner membrane, capsid, and outer membrane are marked by colored backgrounds (IM inner membrane in blue, C capsid in magenta, and OM outer membrane in orange). The coloring scheme corresponds to that in Fig. 1B, C. The 95% confidence interval (N = 14) is indicated by grey shading.

**Fig. S8.**
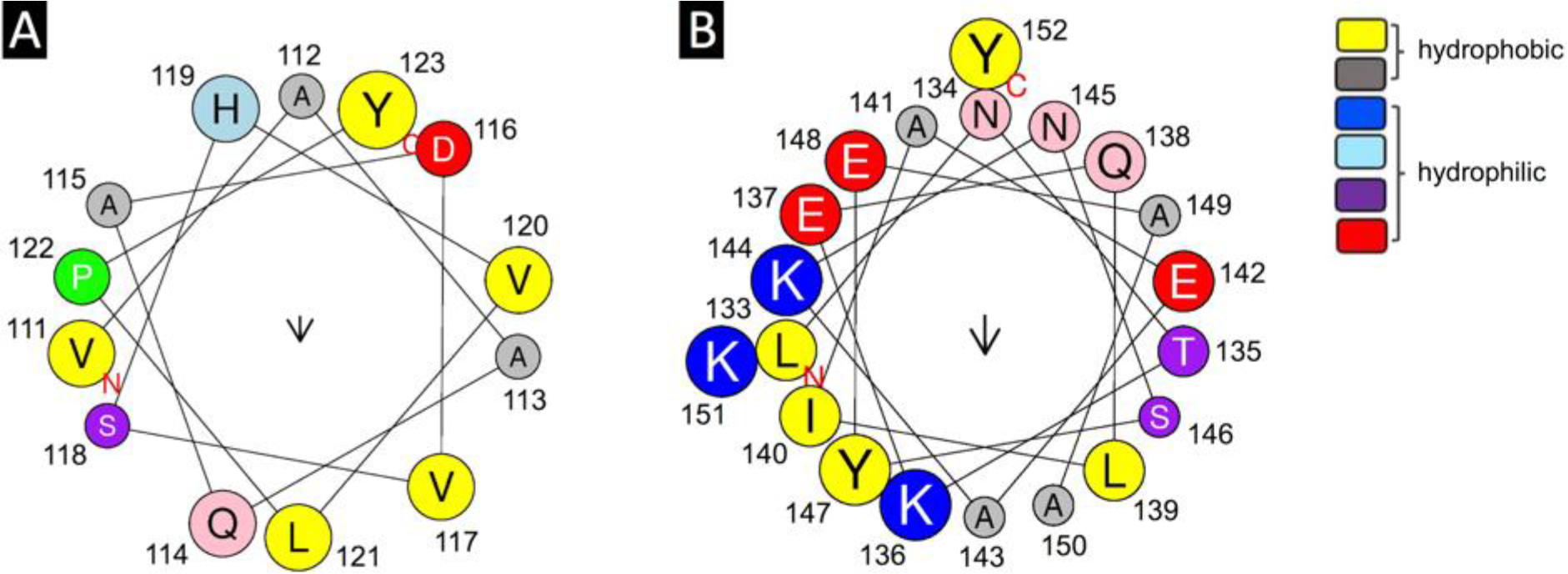
DE loop of J1 domain of EhV-201 major capsid protein contains amphipathic helices α3 and α4. **(A)** Helical wheel representation of helix α3 (residues 111-123) from J1 domain of EhV-201 major capsid protein, prepared using HeliQuest server (*77*), indicating its amphipathic properties. Amino acids with hydrophobic side chains are shown in yellow and grey, and amino acids with hydrophilic side chains are shown in red and blue. The arrow indicates the magnitude and direction of the hydrophobic moment. **(B)** HeliQuest plot of helix α4 (residues 133-152).

**Fig. S9.**
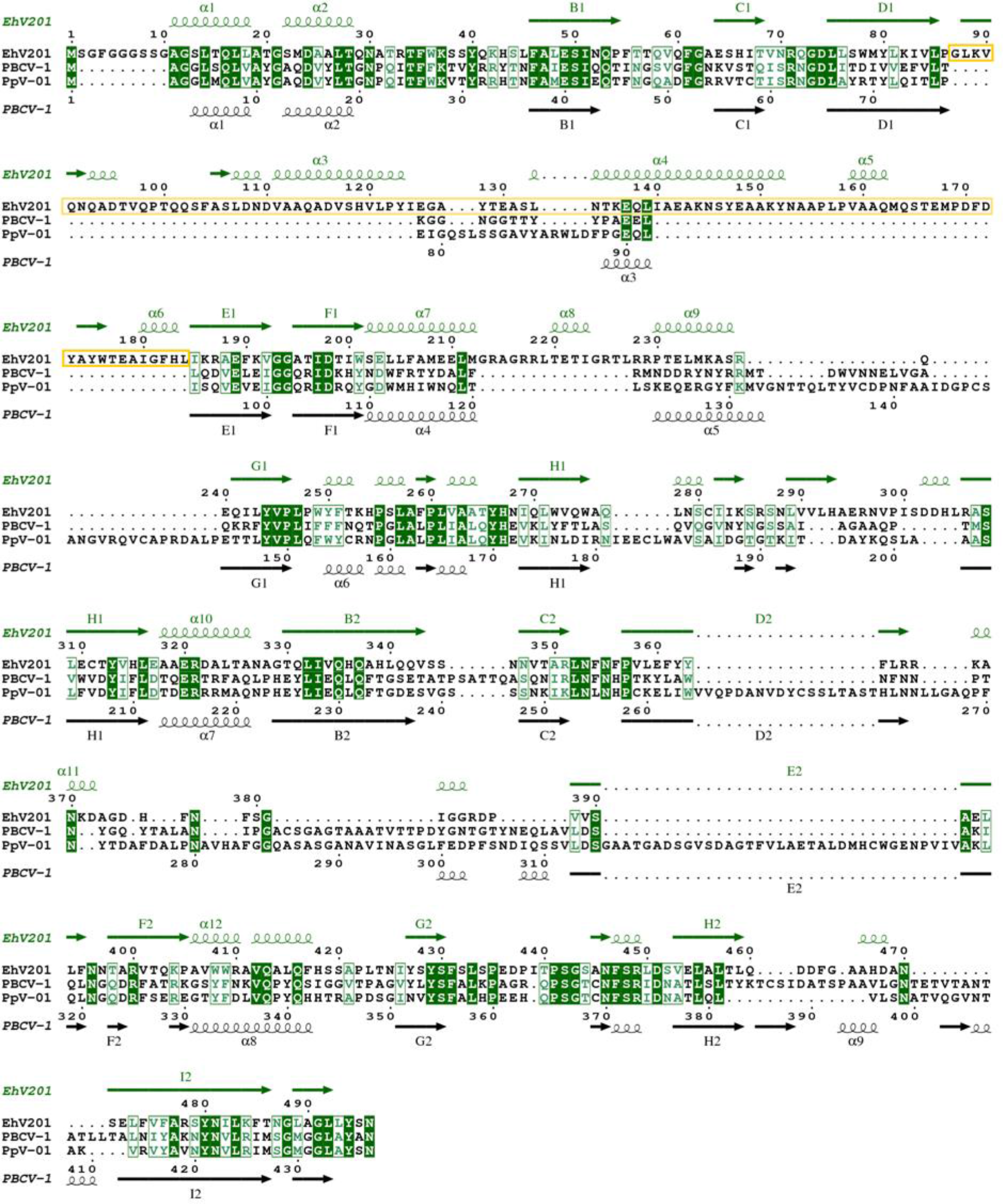
Sequence alignment of major capsid proteins of selected viruses from family *Phycodnaviridae* showing insertions in surface-exposed loops. Major capsid proteins of EhV-201 (Genbank accession code AET97971), PBCV-1 (NP_048787), and *Phaeocystis pochettii* virus (PpV-01, ABU23715) are shown. The 96-residues long insertion in the DE loop of the J1 domain of EhV-201 is highlighted with a yellow box. The insertion is probably unique to coccolithoviruses. Secondary structure elements of the EhV-201 (green) and PBCV-1 (black) major capsid proteins are depicted as arrows for β-strands and spirals for α-helices.

**Fig. S10.**
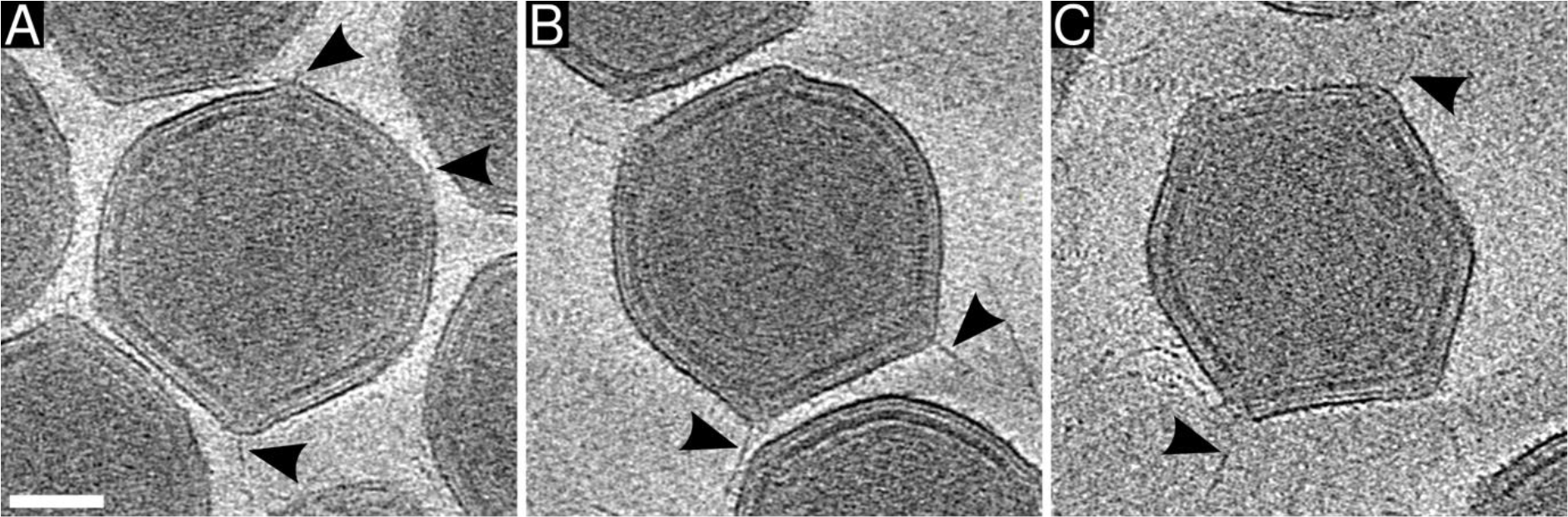
Some vertices of EhV-201 virions are decorated with flexible fibers. **(A-C)** Projection images of 16-nm-thick sections of cryo-tomograms of EhV-201 virions. Fibers attached to some of the virion vertices are indicated by black arrowheads. Scale bar 50 nm.

**Fig. S11.**
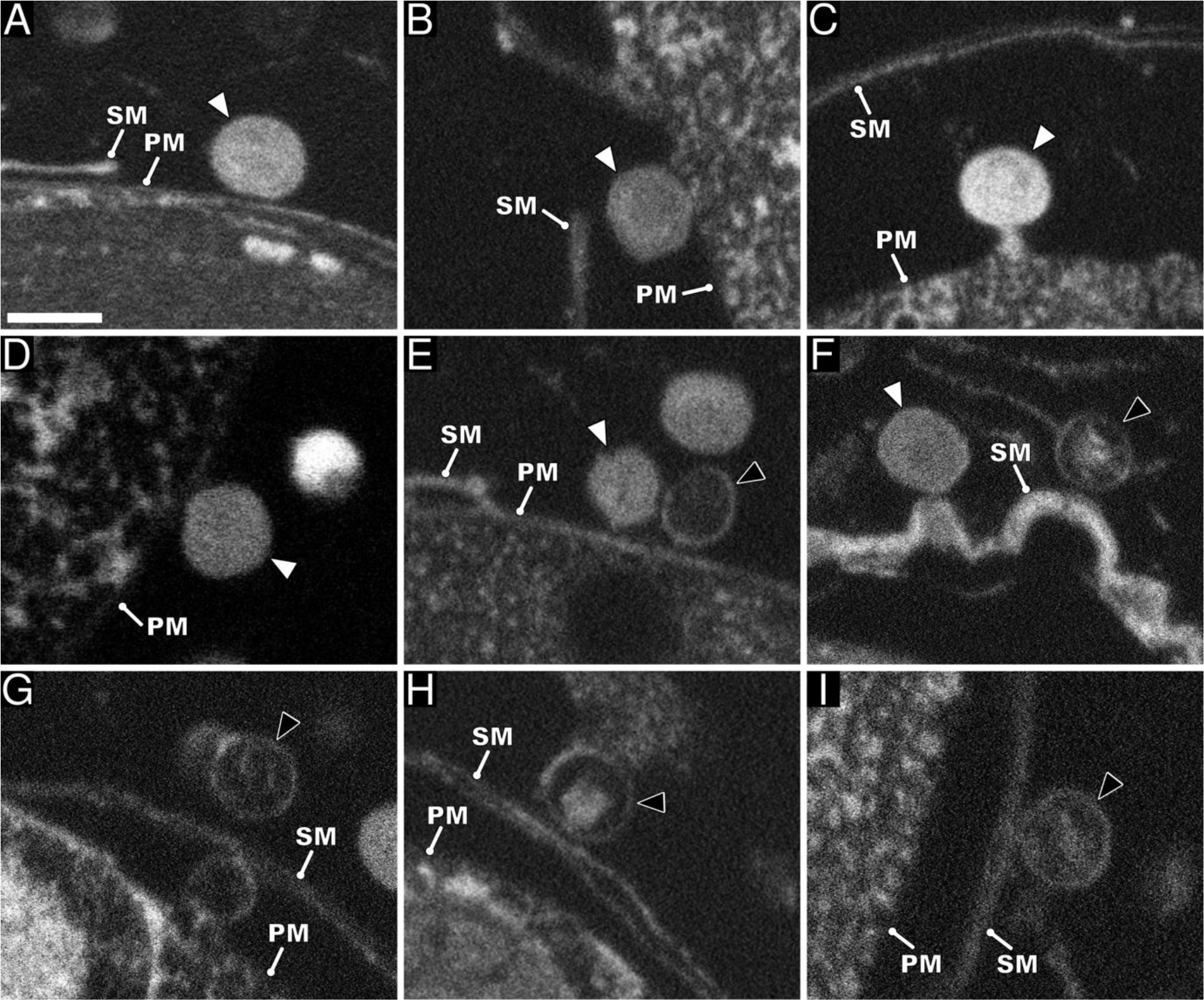
Productive and abortive genome delivery of EhV-201. Scanning electron micrographs of a high-pressure vitrified and resin-embedded sample of *E. huxleyi* cells infected by EhV-201 at MOI = 10, 30 min post-infection. **(A-E)** Productive infection pathway. Genome-containing (A-E) and empty (E) EhV-201 particles attached to plasma membrane. **(F-I)** Abortive infection. Genome-containing (F) and empty (F-I) EhV-201 particles attached to surface membrane of *E. huxleyi* cells. Full particle (white arrowhead), empty particle (black arrowhead with white outline), SM surface membrane, and PM plasma membrane. Please note that the envelope, which covers most *E. huxleyi* cells when imaged using cryo-electron microscopy (Fig. 3), is not resolved in the fixed sample, probably because it was dissolved by the sample fixation procedure or not stained by osmium tetroxide and uranyl acetate used for sample contrasting. Scale bar 200 nm.

**Fig. S12.**
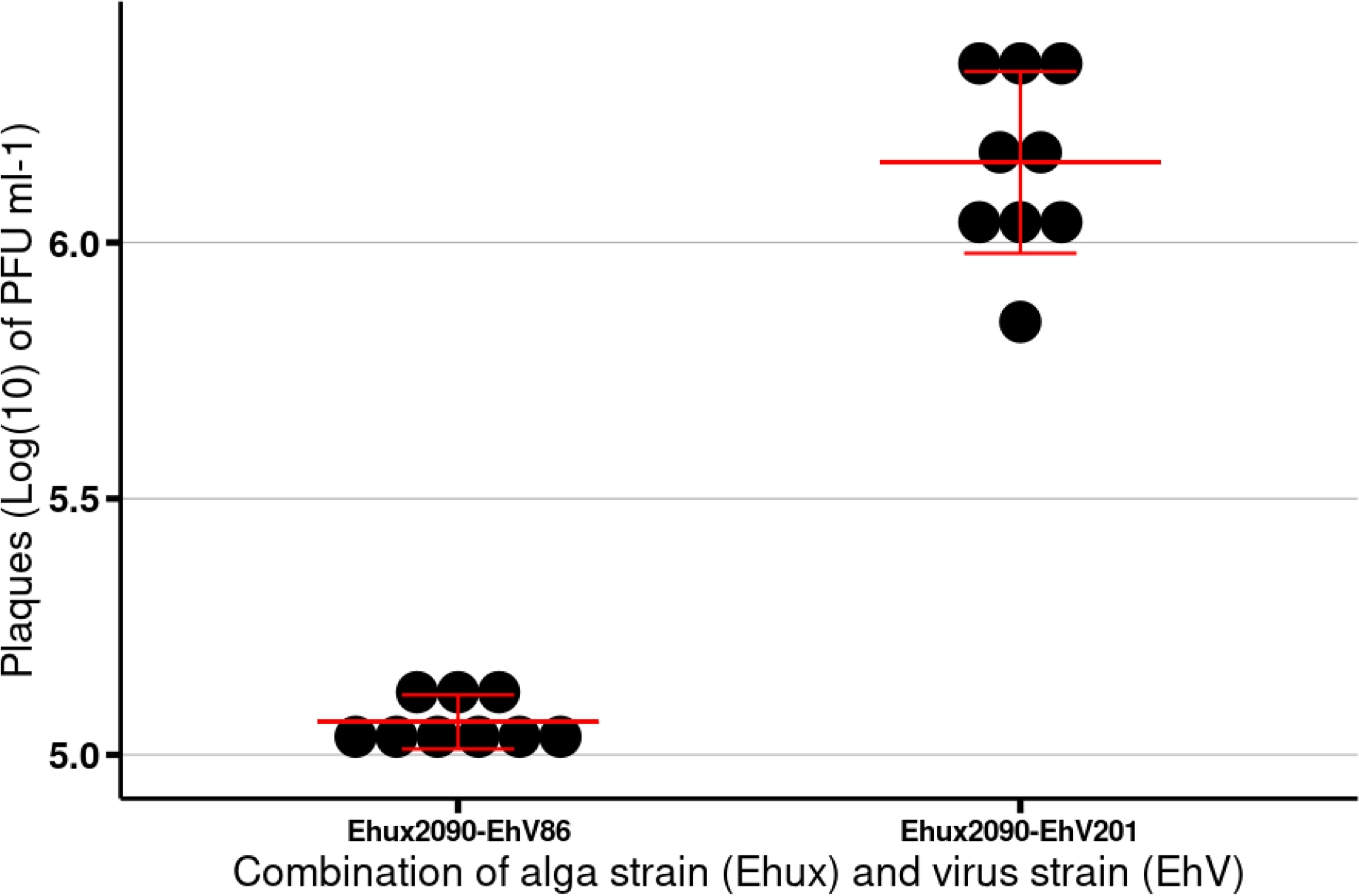
Efficiency of EhV-201 propagation on *E. huxleyi* strain CCMP 2090. Dot plot showing the number of plaque forming units per milliliter obtained for EhV-86 and EhV-201 propagated on *E. huxleyi* CCMP 2090. Mean and standard deviation (error bars) are indicated. N = 3.

**Fig. S13.**
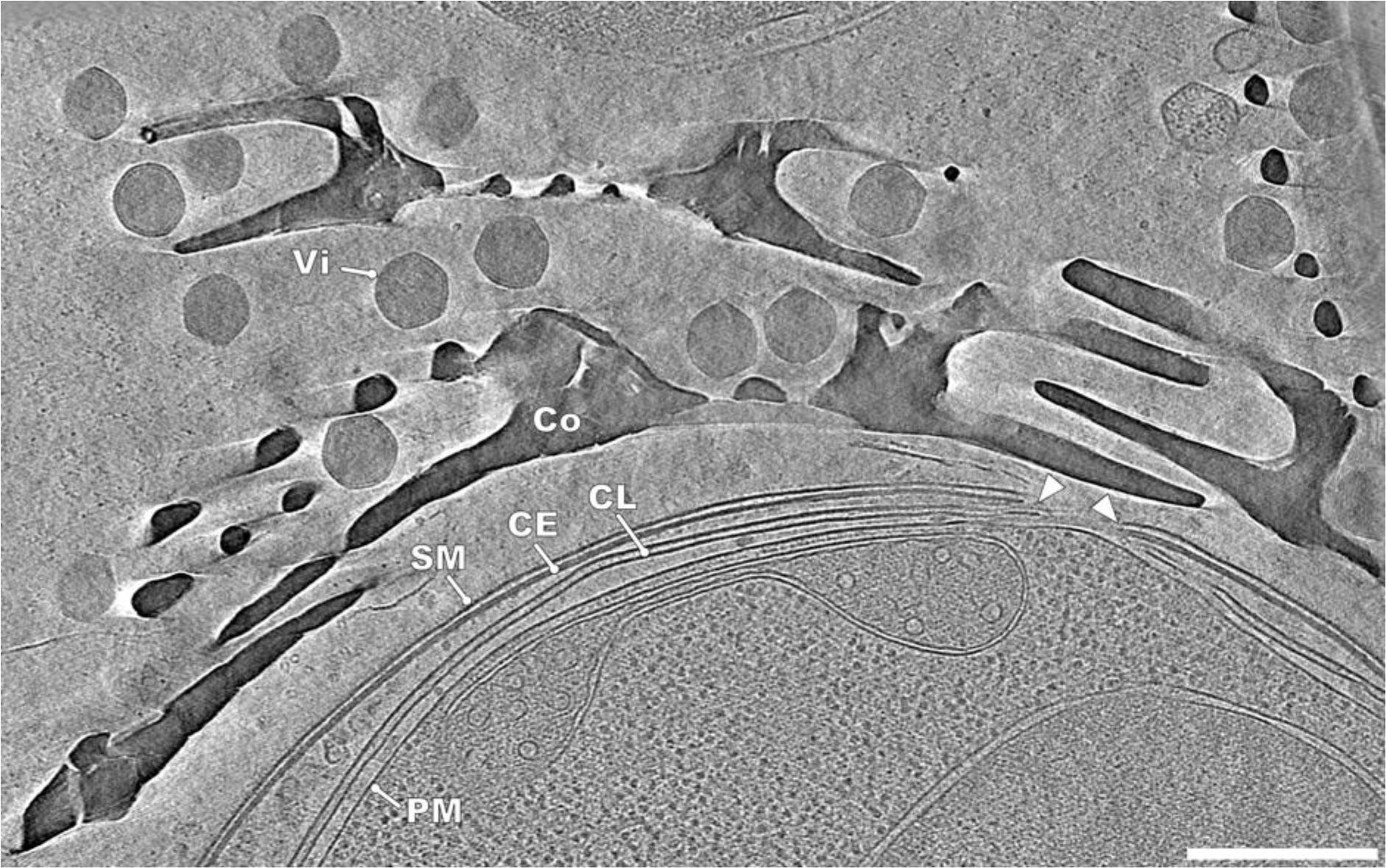
EhV-201 virions can diffuse into *E. huxleyi* coccoliths. A projection image of a 30-nm-thick tomogram section of a cell from the non-calcifying *E. huxleyi* strain CCMP 2090, which spontaneously resumed coccolith production. Co coccolith, SM surface membrane, CE cell envelope, CL cytoplasmic leaflet, and PM plasma membrane. The opening in the surface membrane, cell envelope, and cytoplasmic leaflets is indicated by white arrowheads. The cell is surrounded by a large number of EhV-201 virions (Vi) as it was infected at MOI = 100. Scale bar 500 nm.

**Fig. S14.**
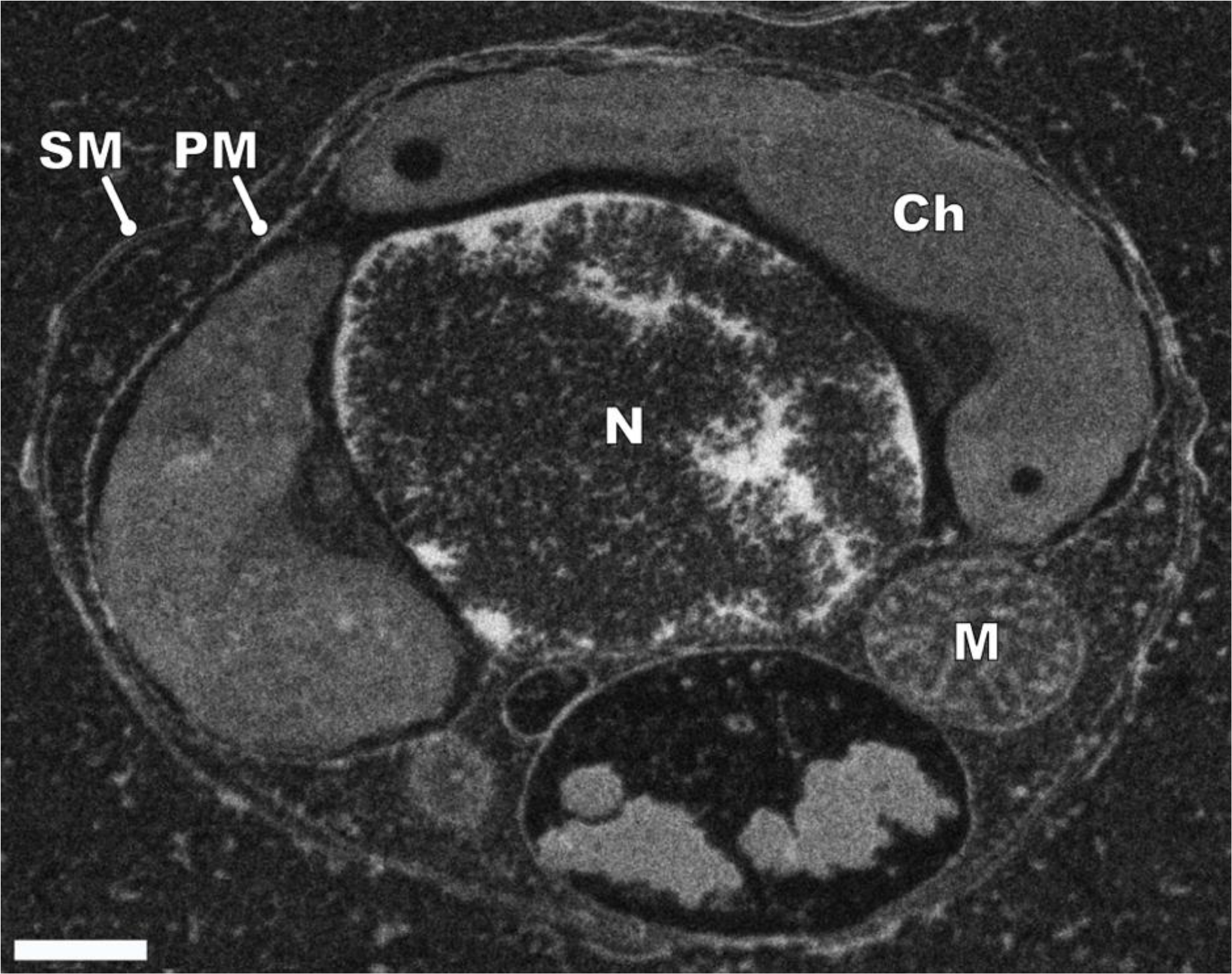
Morphology of native *E. huxleyi* cell. Scanning electron micrograph of a high-pressure vitrified and resin-embedded sample of non-calcifying *E. huxleyi* CCMP 2090 cell. Ch chloroplast, M mitochondrion, N nucleus, SM surface membrane, and PM plasma membrane. Please note that the envelope, which covers most *E. huxleyi* cells when imaged using cryo-electron microscopy (Fig. 3), is not resolved in the fixed sample, probably because it was dissolved by the sample fixation procedure or not stained by the osmium tetroxide and uranyl acetate used for sample contrasting. Scale bar 500 nm.

**Fig. S15.**
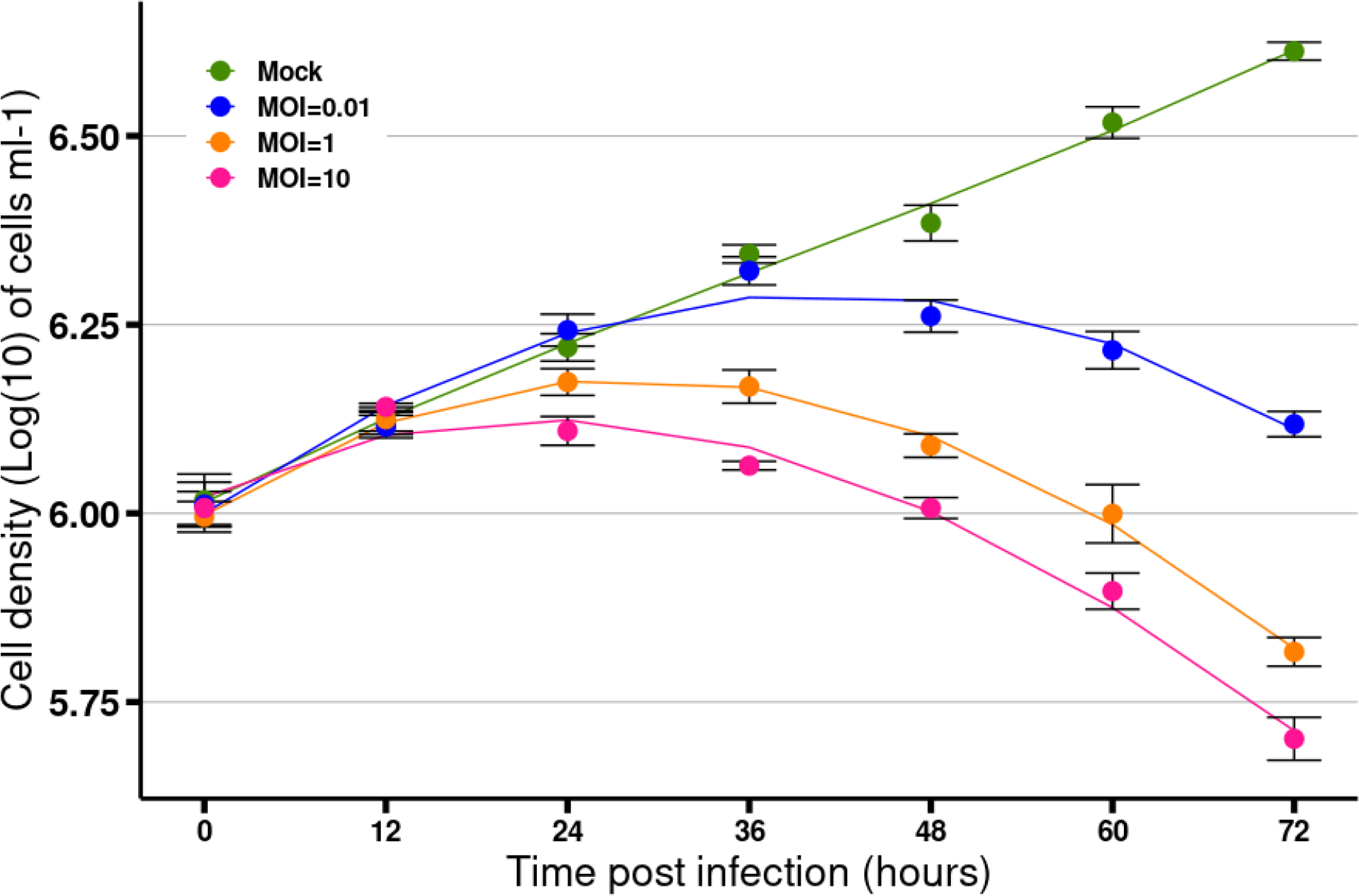
Lysis of *E. huxleyi* culture by EhV-201 at various MOI. Growth curves of *E. huxleyi* CCMP 2090 infected by EhV-201 at MOI 0 (mock), 0.01, 1, and 10. Curves represent the 3rd-order polynomial fit to the data. Error bars correspond to the standard deviation (N = 3).

**Fig. S16.**
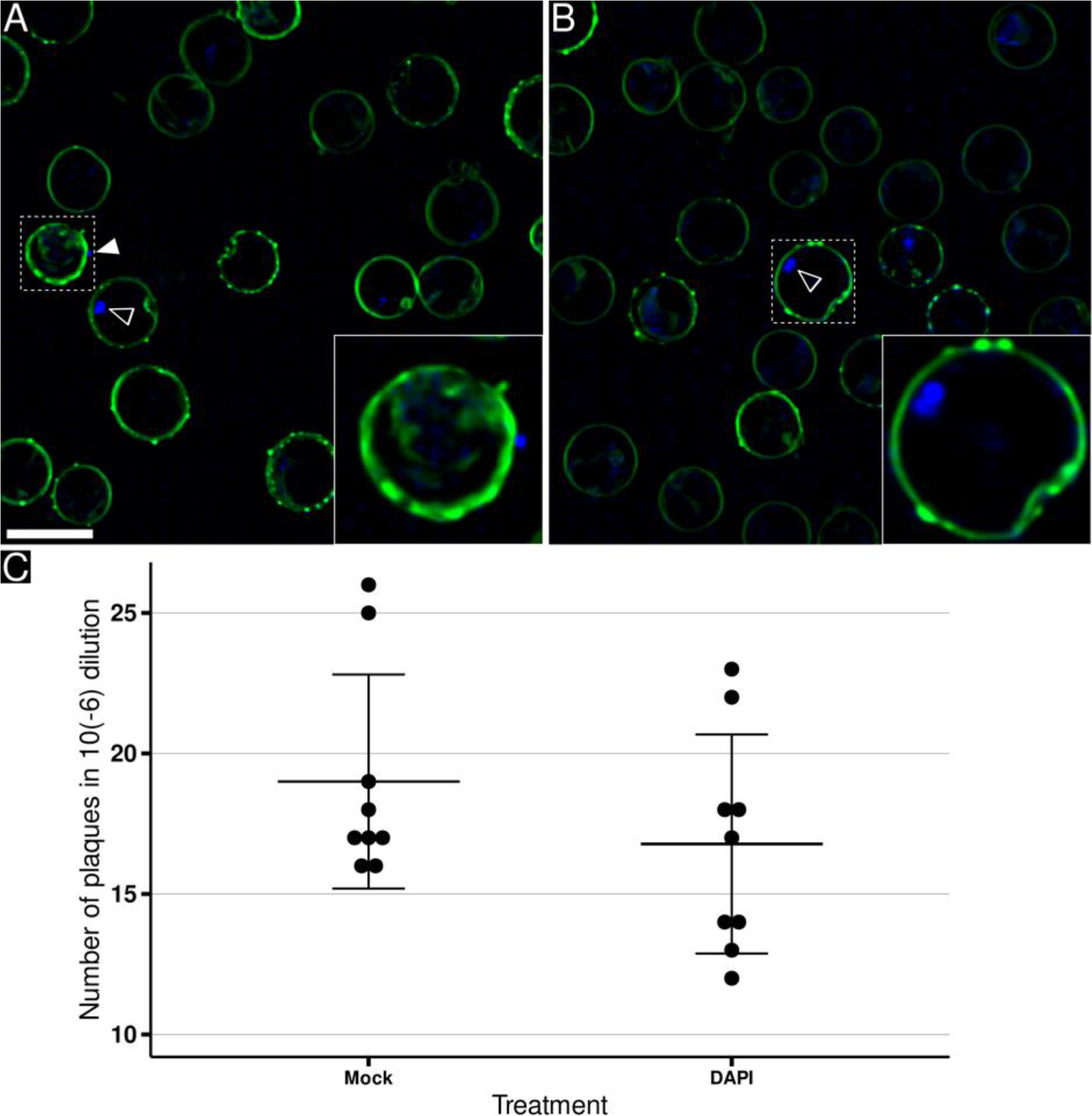
EhV-201 attachment to *E. huxleyi* cells. **(A, B)** Maximum intensity projections of 2,8-μm-thick volumes of fluorescence confocal sections showing plasma membrane of *E. huxleyi* cells in green (stained by FM 1-43) and EhV-201 in blue (stained with DAPI). (A) *E. huxleyi* cells infected at MOI 100. The EhV-201 particle attached to the cell surface is indicated by a white arrowhead. The inset shows details of the cell with a virus attached from the outside. (B) Non-infected control cells. Many *E. huxleyi* cells contain pigment granules that produce a blue signal (indicated by a black arrowhead with a white outline). The inset shows detail of the fluorescent granule inside a control cell. Scale bar 5 µm. **(C)** EhV-201 infectivity is not affected by DAPI fluorescence staining. Dot plot of the number of plaque forming units in 100 µl of a viral lysate with and without DAPI treatment. The mean and standard deviation (error bars) are indicated. The sea water medium-treated group was used as a control (Mock).

**Fig. S17.**
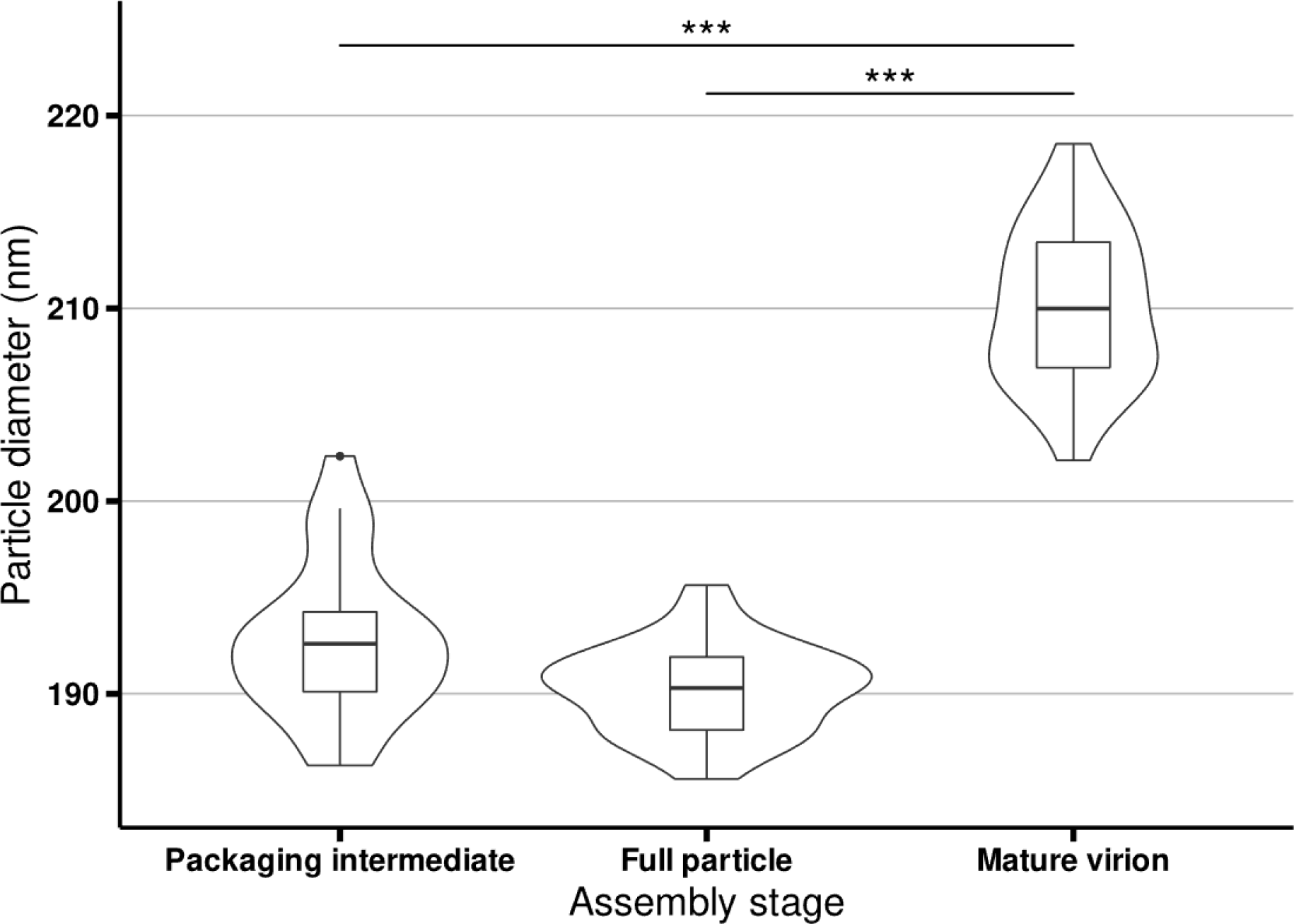
Size distribution EhV-201 assembly intermediates. The maximum outer diameters of genome packaging intermediates, full capsids, and virions were measured from cryo-tomograms of infected cells. Violin plots showing both kernel density and box plot: central black line - median; box - interquartile range; whiskers - 1st and 4th data quartile without outliers; the outlier greater than 1.5 times the interquartile range is depicted by a black dot. N = 25.

**Fig. S18.**
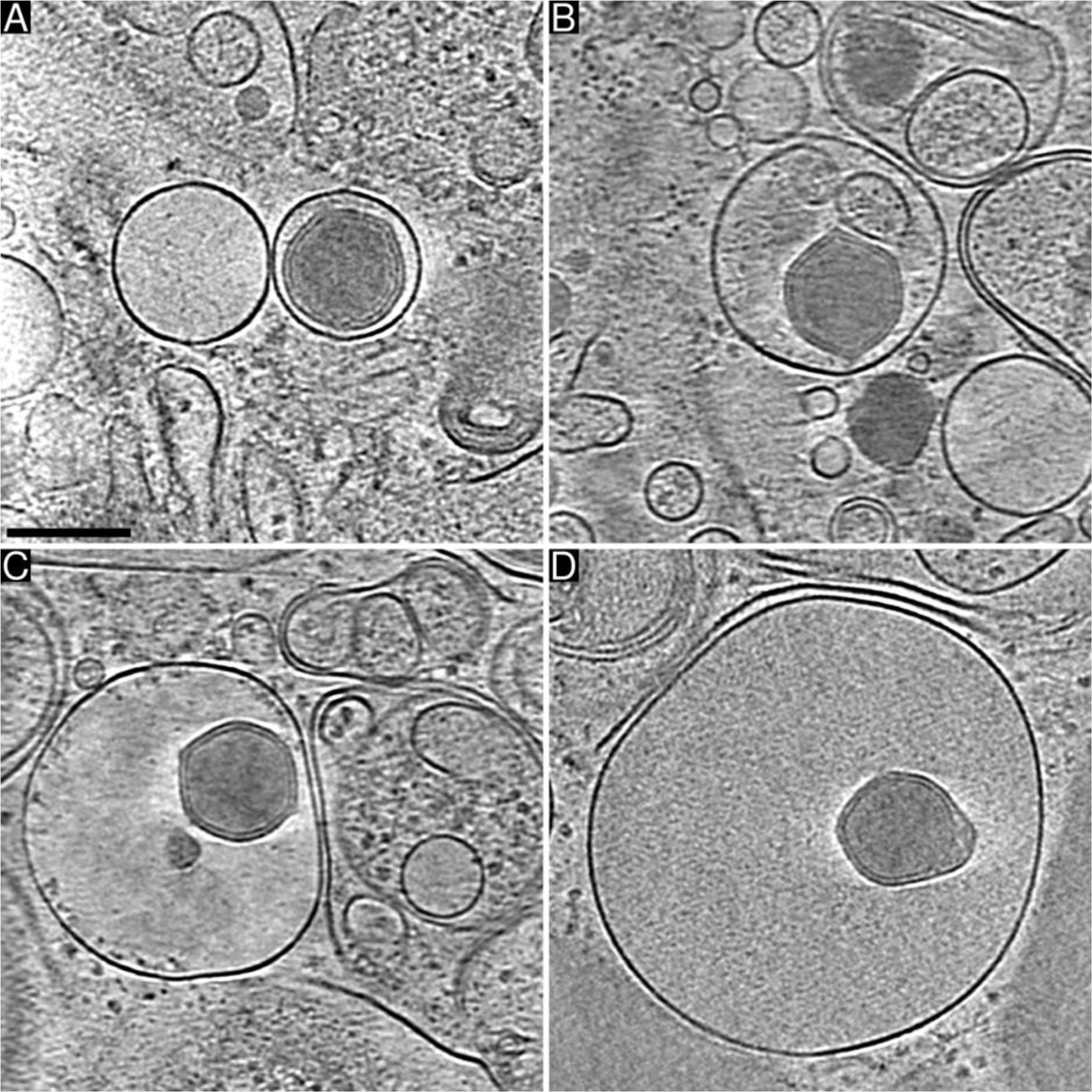
Lysis of infected *E. huxleyi* cells results in release of EhV-201 virions inside vesicles. **(A-D)** Projection images of 30-nm-thick tomogram sections of vesicles released from a lysed *E. huxleyi* cell. Scale bar 200 nm.

**Fig. S19.**
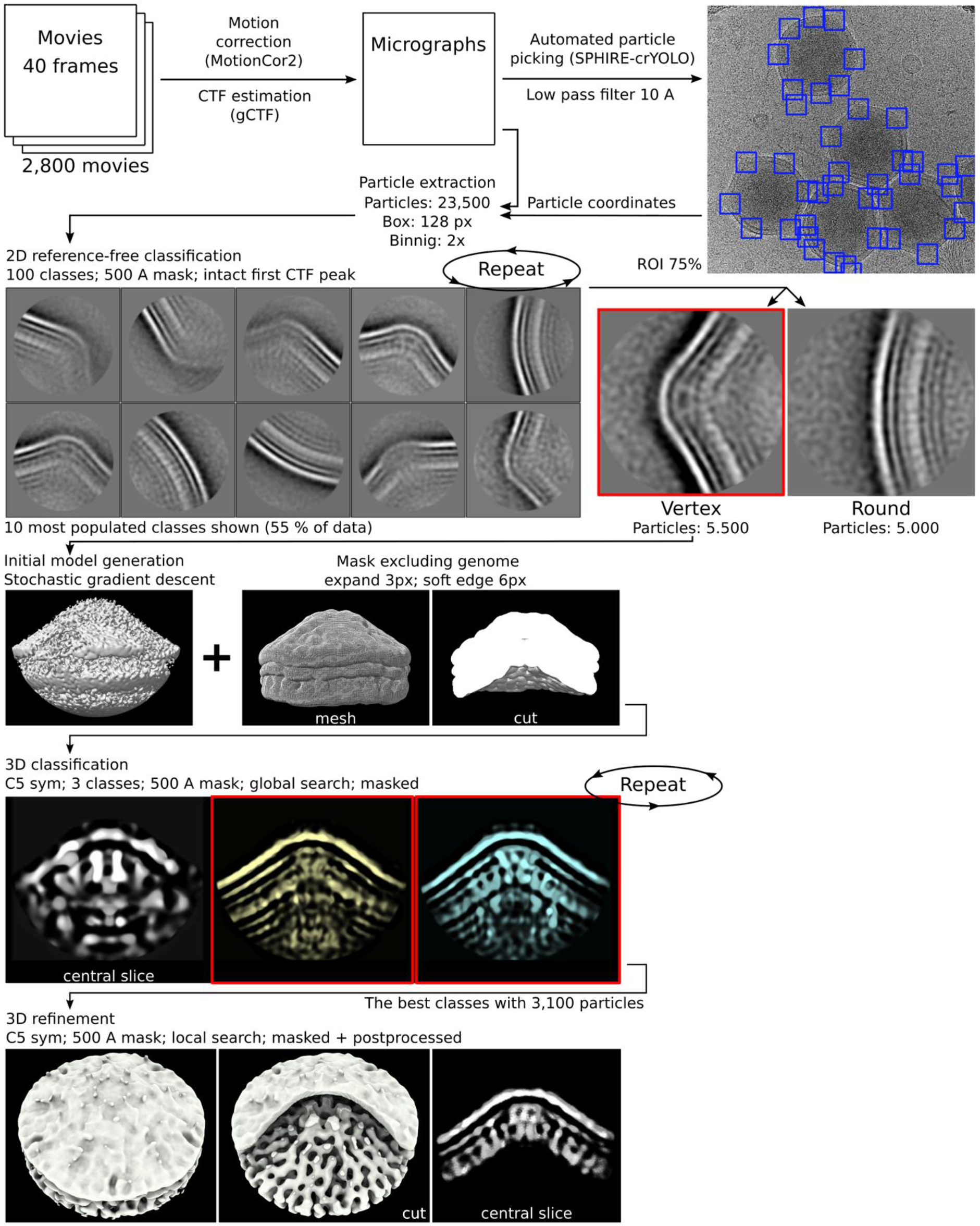
Scheme of single-particle reconstruction of EhV-201 virion vertices.

**Fig. S20.**
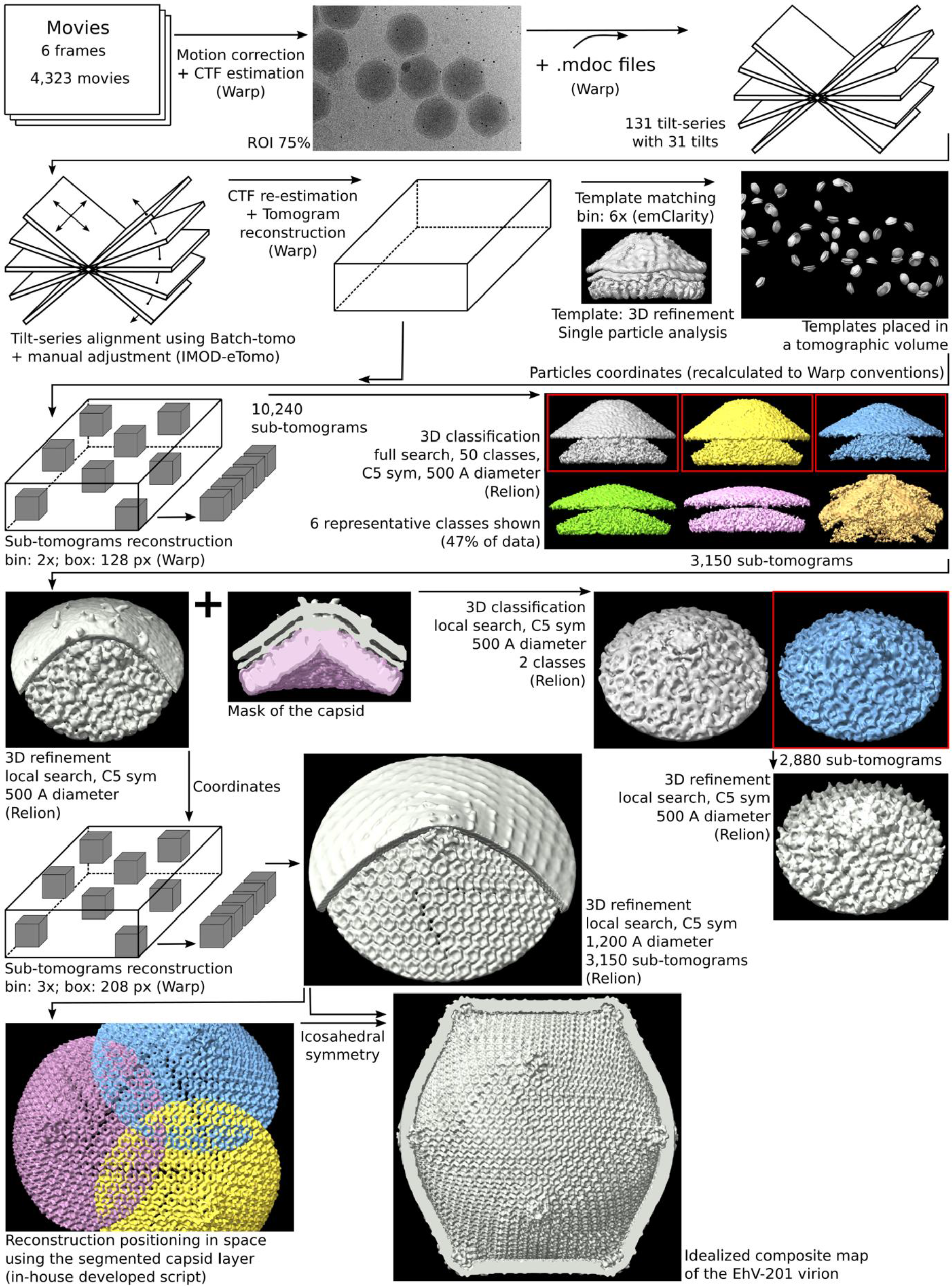
Scheme of sub-tomogram reconstruction of EhV-201 virion vertices.

## Supplementary movies

**Movie S1:**
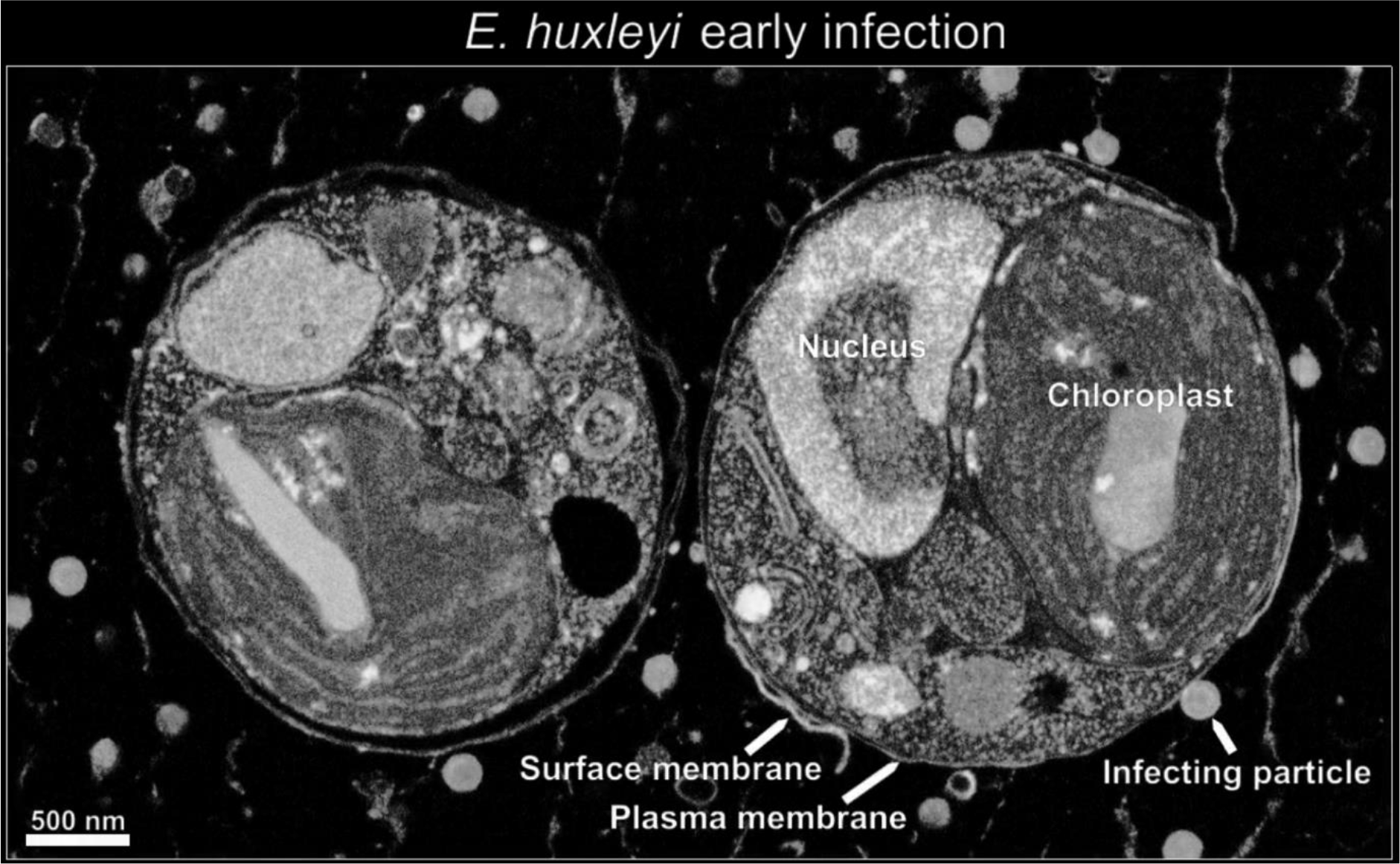
Attachment and genome delivery of EhV-201. The movie shows a sequence of scanning electron micrographs of a high-pressure vitrified and resin-embedded *E. huxleyi* cell infected by EhV-201 at MOI = 10, 30 minutes post-infection.

**Movie S2:**
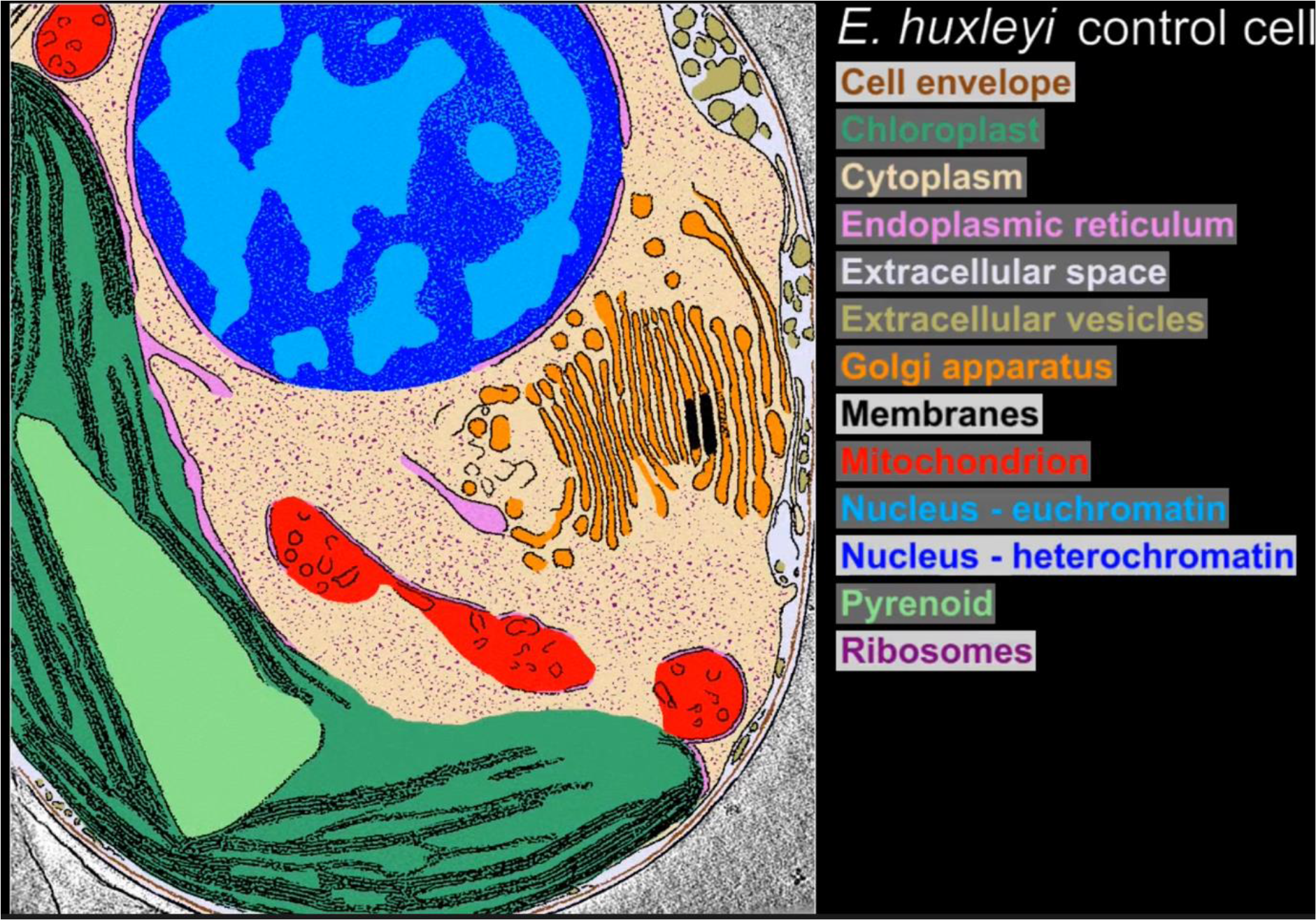
Native structure of *E. huxleyi* cell. The movie shows a series of projection images from a cryo-tomogram of a native *E. huxleyi* cell from the non-calcifying strain CCMP 2090. Scale bar 500 nm. The selected slice is segmented and colored according to organelle type.

**Movie S3:**
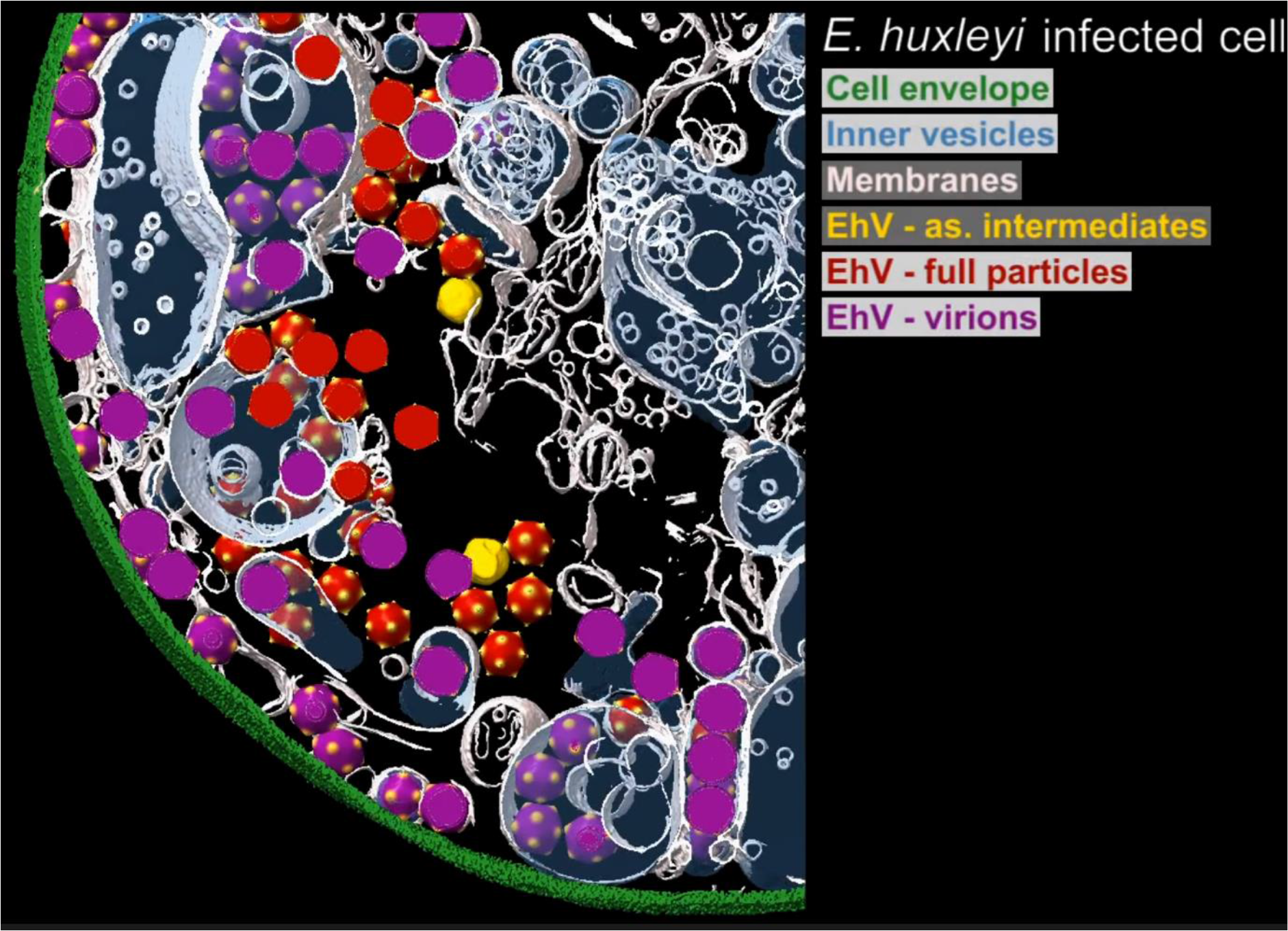
Structure of EhV-201 replication factory. The movie shows a series of projection images and a three-dimensional surface representation of a cryo-tomogram of an EhV-201-infected cell. The cell envelope is shown in green, cellular membranes in white, the content of intracellular vesicles is highlighted with semi-transparent blue, virions in red, full particles in orange, and assembly intermediates in yellow. Scale bar 500 nm.

## Supplementary tables

**Table S1.** Cryo-EM data acquisition parameters, image processing statistics, and structure quality indicators.

| Cryo-EM data collection |  |  |  |  |
| --- | --- | --- | --- | --- |
| Data dedicated for | Single particle analysis | Subtomogram averaging |  | Lamella tomography |
| Microscope settings |  |  |  |  |
| Microscope | Titan Krios G2 | Titan Krios G2 |  | Titan Krios G2 |
| Voltage (kV) | 300 | 300 |  | 300 |
| Projection mode | TEM | EFTEM |  | EFTEM |
| Magnification | 37 k | 42 k |  | 11,5 k to 19,5 k |
| Cs (mm) | 2,70 | 2,70 |  | 2,70 |
| Defocus range (step) (-µm) | 1,2 to 2,4 (0,2) | 2,0 to 4,0 (0,2) |  | 25 |
| Detector | Falcon 3EC | K3 |  | K2 |
| Energy filter | na | Bioquantum |  | Quantum |
| Energy filter mode | na | Zero loss |  | Zero loss |
| Energy slit width (eV) | na | 10 |  | 20 |
| Detector acq. mode | Linear (integration) | Correlated-double sampling (CDS) |  | Linear (integration) |
| Pixel size (Å) | 2,27 | 2,06 |  | 7,4 to 13 |
| Flux on detector (e<sup>-</sup>/px/s) | 72 | 7,5 |  | 50 (*) |
| Dose (e<sup>-</sup>/Å<sup>2</sup>/s) | 14 | 1,61 |  | 0,6 |
| Dose (e<sup>-</sup>/Å<sup>2</sup>/tilt) | na | 2,42 |  | 0,9 |
| Total dose (e<sup>-</sup>/Å<sup>2</sup>) | 28 | 80 |  | 41 |
| Output data format | MRC | TIFF |  | TIFF |
| Number of micrographs | 2800 | 4323 |  | na |
| Number of tilt-series | na | 131 |  | 100 |
| Tilt series setting |  |  |  |  |
| Tilt series acquisition mode | na | Dose symmetric |  | Dose symmetric |
| Starting tilt (deg) | na | 0 |  | 8 (+ or -) |
| Increment step (deg) | na | 3 |  | 3 |
| Maximum tilt (deg) | na | 48 (+ and -) |  | Start tilt +45 and -45 |
| Tilt scheme example (deg) | na | 0;+3;+6;+9;+12;-3;-6;-9;-12;+15;... |  | Start tilt;+3;+6;-3;-6;+9;... |
| Tracking | na | Everytime |  | Everytime |
| Focusing | na | Start tilt + after each 4 |  | Start tilt |
| Cryo-EM reconstruction |  |  |  |  |
| Reconstruction method | Single particle analysis | Subtomogram averaging |  | Tomogram reconstruction |
|  |  | 50 nm diameter | 120 nm diameter |  |
| Initial particle images (no.) | 23500 | 10240 | 3150 | na |
| Particles box | 128 | 128 | 208 | na |
| Binning | 2 | 2 | 3 | 2 to 4 |
| Particle circular mask (Å) | 500 | 500 | 1200 | na |
| Final particle images (no.) | 3100 | 3150 | 3150 | na |
| Symmetry imposed | C5 | C5 | C5 | C1 |
| Map resolution (Å) | 25 | 13 | 18 | 52 to 60 |
| Resolution method | FSC 0,143 (Gold standard) | FSC 0,143 (Gold standard) |  | Theoretical Nyquist |
| Reconstruction software | Relion 3.1 | Relion 4.0 |  | IMOD (eTomo) |
| Accession number | EMD-17650 | EMD-17648,17649<br>PDB 8PFM | EMD-17651 | na |
| Remarks: (*) a' 50 % of electrons absorbed by lamella |  |  |  |  |

**Table S2.** F/2-Si medium composition.

| Alga medium ingredients |  |  |  |
| --- | --- | --- | --- |
| Ingredient | Common name / formula | Manufacturer | Cat.No./Product No./Link |
| Sea water from a functional marine aquaria | Regenerated sea water | Aqua Vala (Brno, Czech Republic) | shop.akvaristika-morska.cz |
| Micronutrients |  |  |  |
| Biotin | Vitamin H or B7 | Sigma Aldrich (Merck) | B4639 |
| Cobalamine | Vitamin B12 | Sigma Aldrich | V6629 |
| Cobalt chloride | CoCl<sub>2</sub> | Sigma Aldrich | 255599 |
| Cupric sulfate | CuSO<sub>4</sub> | Sigma Aldrich | 209198 |
| Ethylenediaminetetraacetic acid disodium salt | Na<sub>2</sub>EDTA | Sigma Aldrich | 324503 |
| Ferric chloride | FeCl<sub>3</sub> | Sigma Aldrich | 236489 |
| Manganese chloride | MnCl<sub>2</sub> | Sigma Aldrich | 221279 |
| Monosodium phosphate | NaH<sub>2</sub>PO<sub>4</sub> | Sigma Aldrich | S0751 |
| Sodium molybdate | Na<sub>2</sub>Mo<sub>4</sub> | Sigma Aldrich | 331058 |
| Sodium nitrate | NaNO<sub>3</sub> | Merck (Darmstadt, Germany) | 106537 |
| Thiamin | Vitamin B1 | Sigma Aldrich | T1270 |
| Zinc sulphate | ZnSO<sub>4</sub> | Sigma Aldrich | 221376 |

**Table S3.** Conditions used to prepare grids with EhV-201 virions for recording data for single-particle reconstruction and tomographic tilt-series and grids with EhV-201-infected *E. huxleyi* cells.

| Cryo-preservation settings |  |  |  |
| --- | --- | --- | --- |
| Data dedicated for | Single particle analysis | Subtomogram averaging | Lamella preparation |
| *Grid specification* |  |  |  |
| Grid material | Copper | Copper | Gold |
| Mesh | 300 | 200 | 200 |
| Coating material | Carbon | Carbon | Gold |
| Hole diameter (µm) | 2 | 2 | 2 |
| Hole spacing (µm) | 1 | 1 | 1 |
| Manufacturer | Quantifoil | Quantifoil | Quantifoil |
| *Glow discharge settings* |  |  |  |
| Device | Gatan Solaris | Gatan Solaris | Gatan Solaris |
| Atmosphere | Hydrogen-oxygen | Hydrogen-oxygen | Hydrogen-oxygen |
| Pirani pressure (Pa) | 0,05 | 0,05 | 0,05 |
| Power (W) | 40 | 40 | 40 |
| Time (s) | 15 | 15 | 15 |
| Glow discharged side | Coating side up | Coating side up | Coating side up |
| *Chamber conditions* |  |  |  |
| Vitrification device | Vitrobot Mark IV | Vitrobot Mark IV | Vitrobot Mark IV |
| Humidity (%) | 100 | 100 | 100 |
| Temperature (°C) | 10 | 10 | 20 |
| Blotting time (s) | 3 | 3 | 12 |
| Blotting force | -2 | -2 | -2 |
| Wait time (s) | 10 | 10 | 10 |
| Blotting paper - front | Filter paper | Filter paper | Wax-soaked filter paper |
| Blotting paper - back | Filter paper | Filter paper | Filter paper |
| Drain time (s) | 0 | 0 | 0 |
| Sample volume (µl) | 3,5 | 3,5 | 4 |
| Vitrification medium | Ethane | Ethane | Ethane |
| Storage medium | Nitrogen | Nitrogen | Nitrogen |

